# Single-cell proteomics workflow for characterizing heterogeneous cell populations in saliva and tear fluid

**DOI:** 10.1101/2025.03.21.644566

**Authors:** Jackson Gabriel Miyamoto, Heloísa Monteiro do Amaral-Prado, Fábio Malta de Sá Patroni, Guilherme Pimentel Telles, Carolina Moretto Carnielli, Guilherme Araújo Câmara, Daniella de Figueiredo, Elaine Cristina Cardoso, Daniela Campos Granato, Alan Roger Santos-Silva, Marcio Ajudarte Lopes, Adriana Franco Paes Leme

## Abstract

Single-cell proteomics (SCP) has advanced considerably but still is largely limited to homogeneous populations and distant from clinical applicability. We present an SCP workflow for assessing the cellular heterogeneity in saliva and tear fluid. Initially, benchmarks were established using a standard HeLa digestion curve, resulting in more than 5,463 protein groups (PGs) at 50 pg. For single HeLa cells, the workflow was improved to minimize contamination and increase quantitative performance, reaching a maximum of 3,785 PGs per single cell. Following, SCP was benchmarked across heterogenous populations of saliva and tear fluid, collected from 10 healthy individuals. By improving cell isolation, contamination control, and DIA-based search and quantitation, single cells from saliva (n=110) and tear fluid (n=149), with average diameters of 8 and 11 µm, respectively, yielded a maximum of 700 PGs per single cell. Downstream analysis indicated overrepresented protein functions, distinct cluster markers and twenty-three validated therapeutic targets identified from single-cell data. Taken together, this study demonstrates the robustness of our SCP workflow applied to biofluids, driving the discovery of biomarkers and therapeutic targets in complex microenvironments.

## Main

Advancements in genomics, transcriptomics, and proteomics approaches have enabled the exploration of complex biological systems, and these technologies are well established at the traditional bulk level for discovering new targets^1^. However, one major disadvantage of studying omics at the bulk level is the scarcity of details about sample heterogeneity and rare cells, with only an average view of molecular expression. Single-cell technologies have been developed to overcome this gap; for example, single-cell transcriptomics (scRNA-seq) is currently a widely used technique with well-established protocols^2^. Although these advances at the RNA level are valuable, it is important to consider that the levels of RNA often do not correlate with the levels of proteins, which serve as the primary functional machinery of cells^3,4^. Single-cell proteomics (SCP) has been an emerging area over the last 10 years and is revolutionizing what we know about cell biology through new advanced liquid-handling robots, enabling the development of one-pot protocols and more sensitive and faster mass spectrometry instruments capable of detecting proteins in low-input and single-cell samples^5–9^.

Biofluids, such as blood, saliva, and tear, are valuable matrices for discovering biomarkers and therapeutic targets, as they can be used to recapitulate disease biology, stratify patients, and monitor responses to therapeutic interventions in diseases^10,11^. They are well recognized as potential “liquid biopsy” sources^10,12–14^. The attractive noninvasive, cost-effective, accessible, and rapid collection features of liquid biopsies can improve patient and professional compliance. Nonetheless, there is still an enormous gap between research advancements and clinical implementation, which is associated not only with the urgent need to advance the validation of discovered signatures in larger cohorts^15^ but also with the lack of sensitivity and specificity required to reliably identify candidate biomarkers and therapeutic targets.

Leveraging SCP approaches for biofluids and other microenvironments can be transformative because of the ability to investigate an individual cell proteome with high cellular heterogeneity, enabling the discovery of health- and disease-related processes and integrating multiple sites. Our group, through connecting the bulk proteome of saliva, blood, and primary and metastatic tumors, revealed head and neck cancer metastasis signatures^13,14^. Thus, investigating biofluids at the single-cell level has the potential to greatly advance the development of a new generation of reliable, mechanism-based health and disease biomarkers and therapeutic targets. Here, we applied an SCP pipeline using single HeLa cells as benchmarks, followed by saliva and tear single-cell analysis from the same healthy individuals. This pipeline integrates single-cell isolation and sample preparation using the cellenONE X1 robot with mass spectrometry analysis on an Orbitrap Astral. Our findings demonstrate both the challenges and feasibility of applying the SCP approach to noninvasive human biofluids for next-generation biomarker and therapeutic target discovery.

## Results

### Establishing a high-throughput proteomic workflow for a standard HeLa digestion curve using Orbitrap Astral mass spectrometer with FAIMS interface

We performed the experiments using a cellenONE X1 robot and a Vanquish Neo UHPLC system coupled with an Orbitrap Astral mass spectrometer (Thermo Scientific, Germany) interfaced with Field Asymmetric Waveform Ion Mobility Spectrometry (FAIMS) Pro (**Figure 1a**). The benchmarks were initially established using a 25 cm × 75 µm ID, 1.7 µm C18 Aurora Ultimate TS analytical column on the Pierce™ HeLa protein digest standard (**Figures 1b-d**) and single HeLa cells (**Figures 1e-g**). An optimized workflow (see methods) featuring a 50 samples-per-day (SPD) active gradient and a sample preparation protocol with 0.05% n-dodecyl-β-D-maltoside (DDM) was implemented. To minimize mass spectrometer contamination, the acquisitions were completed before DDM elution, enhancing system suitability (**Supplementary Figure 1**).

**Figure 1.**
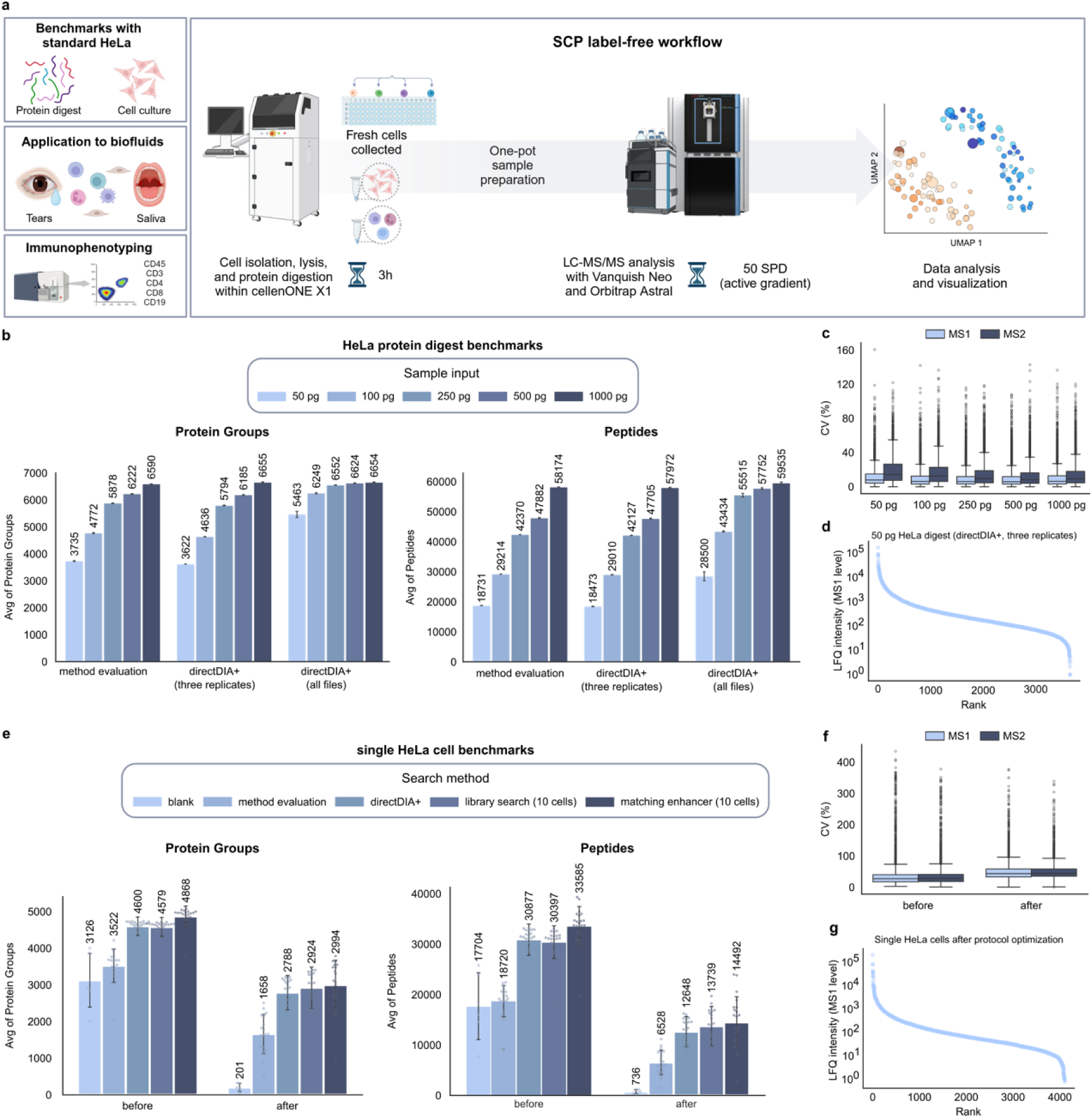
Experimental design and benchmarks for the analysis of HeLa digest, single HeLa cells, and saliva and tear single cells. A) Integrated proteomics workflow for single HeLa cell benchmarks and saliva and tear single cells. B) Average of protein groups or peptides using the Pierce™ HeLa Protein Digest Standard curve (50 pg to 1,000 pg) analyzed using method evaluation and directDIA+ (raw files were treated as replicates of the same condition or all raw files included in a single project, but configured as different conditions) on Spectronaut v19 at the MS1 level. C) Percentage of coefficient of variation (%CV) at the MS1 and MS2 levels of the Pierce™ Standard HeLa Protein Digest curve. D) Dynamic range of quantified protein groups (n=3,652 PG) in 50 pg of HeLa digest at the MS1 level. E) Average of protein groups or peptides for single HeLa cells (n=20) and blank controls (n=6) analyzed via method evaluation, directDIA+, spectral library (10 cells) and DIA-ME on Spectronaut v19, before and after cleaning step optimization of sample preparation within the cellenONE X1 robot. The blank searches were performed independently using directDIA+. F) Percentage of coefficient of variation (%CV) at the MS1 and MS2 levels of single HeLa cells under DIA-ME conditions before and after cleaning step protocol optimization. G) Dynamic range of quantified protein groups (n= 4,106, no imputation) in the single HeLa cells after optimization (DIA-ME) at the MS1 level.

This workflow achieved an average of 3,735 (SD of 13.5) protein groups (PGs) identified in 50 pg of HeLa digest using method evaluation and an average of 5,463 (SD of 115.7) PGs using directDIA+ search on Spectronaut v19 (all files analyzed together) (**Figure 1b**). The estimated quantification of PGs was also evaluated for both MS1 and MS2, with a lower percentage of coefficient of variation (CV) across replicates observed for MS1, as indicated in **Figure 1c**, with medians of 8.05% and 14.53% at MS1 and MS2, respectively. The CVs exhibited minimal fluctuations across the protein standard curve (**Supplementary Table 1**).

Notably, the number of PGs that were both identified and quantified were quite similar (**Supplementary Table 1 and Supplementary Figure 2**a), and the on-the-fly search library increased according to the protein concentration (50, 100, 250, 500 and 1,000 ng), resulting in an increase in the quality of the generated library (**Supplementary Tables 1, 2**). Additionally, PG quantification at the MS1 level of the digest standard demonstrated a dynamic range close to five orders of magnitude (**Figure 1d**, n=3,652 PGs with valid values and no imputation). For HeLa digest quality control, three diagnostic peptides were used as benchmarks to assess the consistency of the standard digest runs (**Supplementary Figure 2**b).

The quality of the single-cell one-pot protocol for HeLa cells was assessed by evaluating two trypsin autolysis peaks, VATVSLPR, m/z 421.7584, +2, and LSSPATLNSR, m/z 523.2855, +2, across all single-cell acquisitions. These peptides demonstrated excellent reproducibility, with a retention time standard deviation of only 3–4 seconds for VATVSLPR (**Supplementary Figure 3**a). Among the identified stripped sequences in single cells, 71% were doubly charged, whereas 25% were triply charged, with a median peak width of 5.3 seconds. Additionally, 84% of the sequences exhibited no missed cleavage sites, and 14% presented a single missed cleavage site (**Supplementary Table 1**). These findings suggest highly efficient single-cell protein digestion and effective filtering of charged species by the FAIMS device, set to a CV of -48 V.

### Defining controls and quantitative performance for single HeLa cell analysis

In parallel, optimizations in the protocol for single-cell isolation were carried out under different conditions for the controls, which were referred to as blanks. We evaluated three types of blanks for the single-cell experiments: a) Blank 1, which refers to wells containing only the master mix dispensed with no cells isolated (n=12 wells for saliva and tear; n=6 wells for HeLa cells) and prepared in the same 384 plates where the cells were isolated to assess cross-contamination within the workflow; b) Blank 2, which includes wells with master mix (n=10 wells) prepared in a plate without isolated cells on the day following the cell isolation to evaluate baseline contamination; and c) Blank 3, which includes 10 LC‒MS/MS runs without injection and serves as an analytical column carryover control. Blanks 2 and 3 were used exclusively for saliva and tear experiments.

We carried out independent searches for the blanks using directDIA+ on Spectronaut v19, with the same parameters as those applied for the single-cell searches. Initially, a mean of 3,126 (SD of 731) PGs and 17,704 (SD of 6,043) peptides were identified in the blank controls (blank 1, n=6) (**Figure 1e**, before optimization). To minimize this high protein identification in blank 1, we implemented additional cleaning steps, before and after cell isolation, for the medium-sized PDC (Cellenion Piezo Dispense Capillary) and MS-grade water on the system bottle (see methods). To evaluate the system after cleaning step optimization, blank 1 (n=6) was added to 384-well plates during single HeLa cell isolation. The result was a striking decrease in identifications, with a mean of 201 PGs (SD of 109) and 736 peptides (SD of 379) (all 6 files combined, directDIA+, MS1 level, default Qvalue cutoffs) (**Figure 1e**).

The exclusive PGs identified in each blank before and after optimization were investigated, and the top enriched GO biological process was intermediate filament organization (GO:0045109; adjusted p-value<0.05), which includes keratin proteins and still remained after protocol optimization (Venn diagrams in **Supplementary Figures 3e, f** and **Tables 3, 4**). Additionally, contamination was also evaluated by linear regression analysis of single-cell diameters and PGs before and after cleaning, revealing a significantly greater R-squared values after optimization than before (p<0.05, R² = 0.42 and 0.034, respectively) (**Supplementary Figures 3h, i**).

For single HeLa cell benchmark identifications, two additional search strategies were employed: generating a spectral library using Pulsar from a bulk of 10 isolated cells or co-analyzing these files with cell samples as high-input (matching enhancers or DIA-ME), as described by Krull et al.^16^ (**Figure 1e**). Prior to optimization, no significant improvement in proteome coverage was observed using the library search compared to directDIA+. However, a 5% increase in proteome coverage was achieved with the DIA-ME strategy (**Figure 1e**). After optimization, the DIA-ME search method still yielded superior identifications of PGs and peptides, although the difference compared to the spectral library search was less pronounced, with a 2% increase (**Figure 1e**). The CVs for single HeLa cells at the MS1 level had medians of 27.73% and 44% before and after optimization, respectively, whereas at MS2, the medians were 28.45% and 45%, respectively (**Figure 1f**). The increase in the CV values after optimization suggested that background contamination was evenly distributed during sample preparation, contributing to the lower CVs. Additionally, MS1 showed a slight increase in the number of data points, indicating improved quantitation, with a dynamic range approaching five orders of magnitude (n=20, no imputation). This performance is comparable to that of 50 pg of HeLa digest, which, while identifying nearly twice as many PGs and peptides, demonstrated a similar dynamic range (**Figure 1g**).

### Establishing strategies for cell isolation, contamination control and data analysis for saliva and tear single-cell proteomics

The establishment of HeLa benchmarks at the bulk (total protein digest) and single-cell levels demonstrated the robustness and feasibility of this pipeline for SCP. It improved the quality of the isolated cells and minimized contamination, as shown by the analysis of blank 1 before and after protocol optimization.

Single-cell isolation was performed in saliva and tear fluid (initially, 200 cells were isolated from the saliva and tear fluid). To minimize cross-contamination from remaining HeLa cells, an exclusive and cleaned PDC for human biofluids was employed. The samples were freshly collected from a pool of ten healthy individuals, and morphometric analysis revealed high heterogeneity in the cell sizes obtained during the isolation of both biofluids: 5– 45 µm in diameter for saliva and 5–20 µm in diameter for tear fluid, with an average of 11 µm in diameter for saliva and 8 µm in isolation for tear fluid (see methods and **Supplementary Figures 4a,d and 5a,d**). The morphometric parameters for single cells were obtained from cellenONE X1 and are described in **Supplementary Table 5**. Notably, characterizing the proteome of single cells from saliva and tear fluid is challenging at various stages, from isolation to data analysis, as clinical samples have a greater diversity of cell types and sizes than do immortalized cell culture lines, such as HeLa^17^ and HEK293^18^, or even more homogeneous cell populations previously classified by fluorescence-activated cell sorting (FACS) (**Supplementary Figures 3g, 4d, 5d**).

One of the main challenges encountered with these diverse samples was setting up the most suitable data analysis strategy for the generated MS files. The MS data for single-cell saliva and tear fluid were processed separately on Spectronaut, enabling a DIA-ME search. However, some files did not meet the global protein cutoff of 0.01, even though the run-level protein FDR threshold was set to 0.05. This discrepancy is likely attributed to the inherent cellular heterogeneity and to the low intensity of MS/MS spectra typically observed in single-cell analyses. These effects were especially pronounced in saliva cells, which exhibited greater variability in cell size than those observed in the tear fluid data (**Supplementary Figures 4a, 5a**). To address this challenge, two strategies were implemented: first, the global protein FDR threshold (experiment wise cutoff) was raised to 0.05 for saliva samples; second, raw files with insufficient peptide-spectrum matches (PSMs) to accurately determine Cscore cutoffs were excluded, and the dataset reprocessed using the default threshold of 0.01. As a result, the ratio of the precursor hits of the decoy database to total hits (decoy + database) decreased from 0.44 and 0.48 to 0.39 and 0.45 for the saliva- and tear-filtered data, respectively, as illustrated in **Supplementary Figures 6a-b**.

After applying the filtering strategies, the searches were repeated with the default cutoffs, resulting in 110 saliva and 149 tear single-cell raw files, which led to an improved on-the-fly spectral library and an increased number of identified PGs from 616 to 1,101 for saliva and from 899 to 1,122 for tear fluid (**Figures 2a, b** and **Supplementary Tables 6, 7**). The spectral library recoveries were 55.3% and 52.3%, with data completeness rates among the PG profiles of 32.6% and 37.2% for saliva and tear single cells, respectively (**Supplementary Figures 4e, 5e**). Notably, 60% of the saliva and 98% of the tear cells excluded from the analysis were smaller than 11 µm in diameter, likely reflecting low proteome coverage in these smaller cells. Additionally, morphometric analysis revealed a greater CV in the diameter of saliva cells (**Supplementary Table 5**) than in that of tear cells (50.8% vs. 28.5%), highlighting greater cellular heterogeneity in saliva compared to tear cells. Krull et al.^16^ reported a decrease in protein identification with large signal intensity differences among samples, which may explain the increase in PGs after filtering the files. Differences in the total ion chromatograms and profiles of representative raw files between nonexcluded (**Figures 2c, e**) and excluded samples (**Figures 2d, f**) were evident.

**Figure 2.**
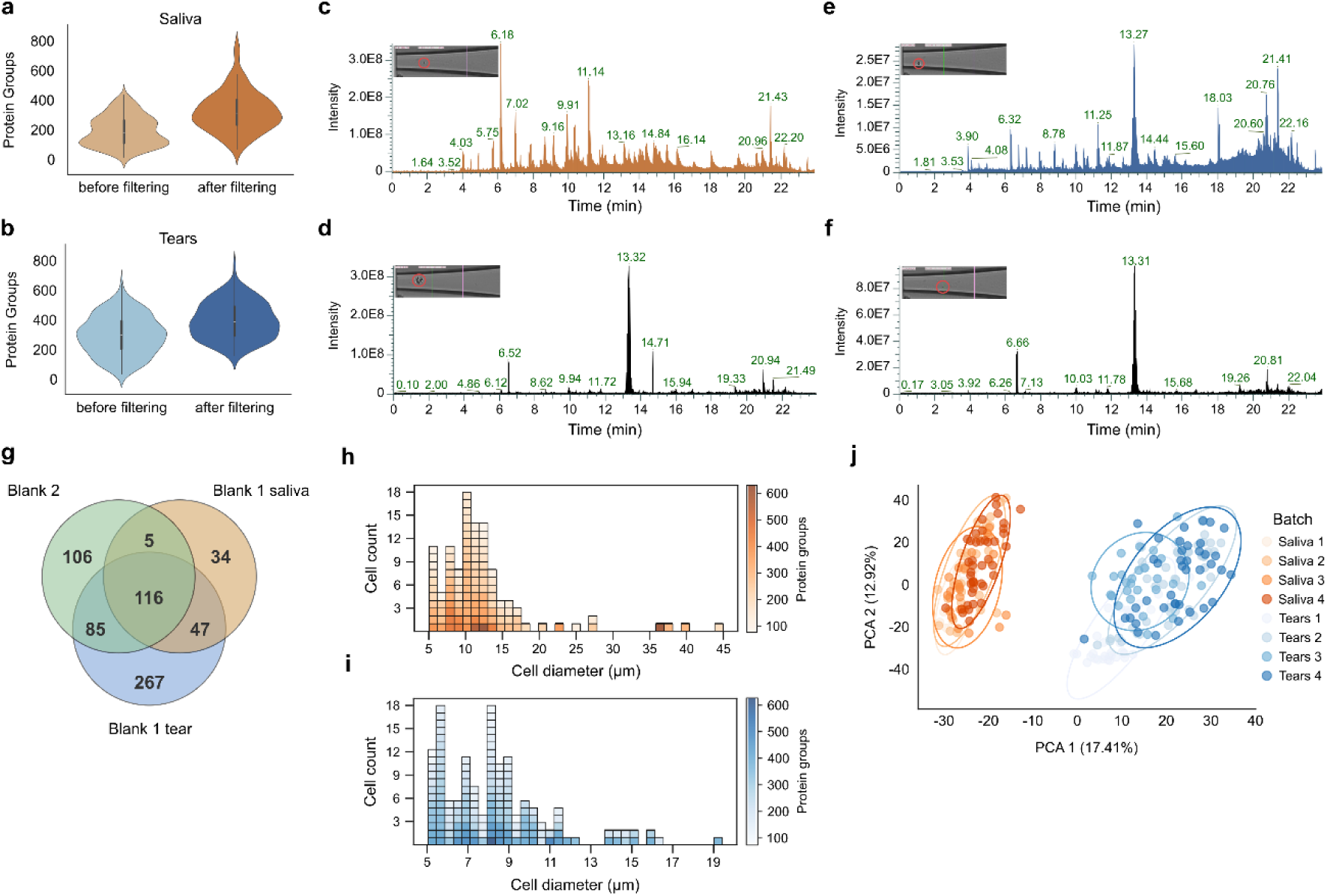
Optimizing parameters for saliva and tear single-cell analysis. A, B) Violin plots showing the distribution of PGs identified in saliva (orange) and tear (blue) single-cell samples. Unfiltered files identifications were accessed by raising the experiment protein Qvalue cutoff to 0.05. Filtered files were processed using the default search parameters. C, D, E and F) Representative chromatograms and photographs of nonexcluded (upper panels) and excluded (lower panels) samples for saliva (C and D) and tear (E and F) single cells. G) Venn diagram of blank 1 (saliva and tear), carried out in the respective 384-well plates of saliva and tear single cells; and blank 2, carried out on the following day of the cell isolation, without any cells in the experiment. H, I) Histograms of cell diameters on the basis of the number of PGs for saliva (H) and tear (I) single cells. J) Principal component analysis (PCA) of the batch effect through LC‒ MS/MS runs; each dot represents a cell. The analysis was conducted in two stages: an initial search using on-the-fly spectral libraries followed by the construction of a communal library, after which the data were reanalyzed and externally normalized.

Regarding the blanks, the blank 3 contained an average of twenty-seven PGs (**Supplementary Table 8**, including cRAP sequences), with the most abundant identifications representing a negligible 1% carryover, which was calculated by the average intensity of the most abundant PGs identified in blank 3 divided by the average MS1 absolute quantity in filtered single cells of both biofluids. For carry-over calculations, it is not easy to define the best strategy considering the variations in size and protein content. In contrast, for blanks 1 (**Supplementary Tables 9, 10**) and 2 (**Supplementary Table 11**), the 116 shared PGs, which are related mainly to the biological process of intermediate filament organization (GO:0045109, adjusted p-value<0.05), were removed from the filtered data to eliminate background identifications and facilitate a more accurate exploration of the heterogeneity of single cells (**Figure 2g** and **Supplementary Tables 12, 13**). The batch effects were evaluated using a Principal Component Analysis (PCA) plot for all filtered data, which was searched against a communal library built from independent searches of saliva and tear samples (see methods). Although the protein digestion of saliva and tear single cells was performed on independent days, the LC‒MS/MS runs were randomized across the saliva and tear single-cell samples (**Supplementary Table 14**). The PCA plot revealed two distinct clusters, representing saliva and tear samples, highlighting the clear separation between the two biofluids (**Figure 2j**). However, within each biofluid cluster, there was an overlap between batches, indicating negligible batch effects within the saliva and tear single-cell samples (**Figure 2j** and **Supplementary Figures 4c, 5c**).

We further demonstrated that there are differences in the distribution of cell diameters, particularly in range, number of PGs per cell, and counts of cells of a specific diameter. Single saliva cells spanned a broader range of cell diameters, extending up to 45 µm, with a majority concentrated between 8 and 14 µm and greater than 20 µm (**Figure 2h** and **Supplementary Figure 4**a). In contrast, tear cells showed a narrower range, up to 19 µm, with higher counts occurring at smaller diameters, particularly approximately 5–8 µm, and a steeper decrease after 8 µm (**Figure 2i** and **Supplementary Figure 5**a). Representative isolated cells and their diameters are shown in **Supplementary Figures 4d and 5d**. In addition, there was no clear relationship between cell size and the number of PGs identified (**Figures 2h, i** and **Supplementary Figures 4b, 5b**).

To complement the characterization of the cells from saliva and tear fluid, targeted immunophenotyping was performed using the same biofluid samples utilized for single-cell isolation. The cells were stained with immune surface markers, including anti-CD45, anti-CD3, anti-CD4, anti-CD8, and/or anti-CD19 antibodies. The results confirmed the presence of both T and B cells in the saliva and tear samples. In saliva, 51.9% of the cells were viable, and 76.3% of these viable cells were identified as CD3^+^ T cells. Within the CD3^+^ T-cell population, 8.07% were CD4^+^ helper T cells, and 2.58% were CD8^+^ cytotoxic T cells, indicating that a small proportion of these T-cell subsets were present in the sample. B cells were identified as CD19^+^ in 5.18% of the CD3^-^ T population. In tear samples, 71.6% of the cells were viable, of which 4.42% were leukocytes, as identified by CD45 staining. Among these leukocytes, 56% were CD3^+^ T cells. Within the CD3^+^ T-cell subset, CD4^+^ T cells were predominant (91.6%), whereas CD8^+^ T cells were predominant (10.7%). B cells were also detected in tear samples, with 51.5% of CD3^-^ T cells expressing CD19 (**Supplementary Figures 7, 8**). Among the surface markers, only CD45 (P08575) was identified in 6% of the total tear cells analyzed via LC‒MS/MS, as represented by Uniform Manifold Approximation and Projection (UMAP) results, indicating the presence of a few colored cells, similar to the flow cytometry results (4.42% of cells stained with anti-CD45) (**Supplementary Figure 8**i).

### Downstream functional analysis of heterogeneous single-cell populations from biofluids

Following single-cell metric analysis of saliva and tear fluid samples, we investigated the biological significance of the identified proteome. First, two heatmaps, representing the tear and saliva datasets, highlighted distinct clustering and PG distribution patterns. The saliva heatmap, which is based on the Spearman-Ward method, covered 985 PGs and exhibited a more balanced cluster distribution (Cluster 1 with 57 cells and Cluster 2 with 53 cells). The saliva data also revealed more localized and intense PG expression clusters than the more evenly distributed patterns observed in the tear heatmap (**Figure 3a**). In contrast, the single-cell tear heatmap, generated using the Kendall-Ward method, included 1,005 proteins and separated the cells into two main clusters: Cluster 1 with 30 cells and Cluster 2 with 119 cells. It also displayed broader and more diffuse PG expression patterns across cells (**Figure 3b**). These differences may reflect variations in single-cell complexity, size and PG expression variability.

**Figure 3.**
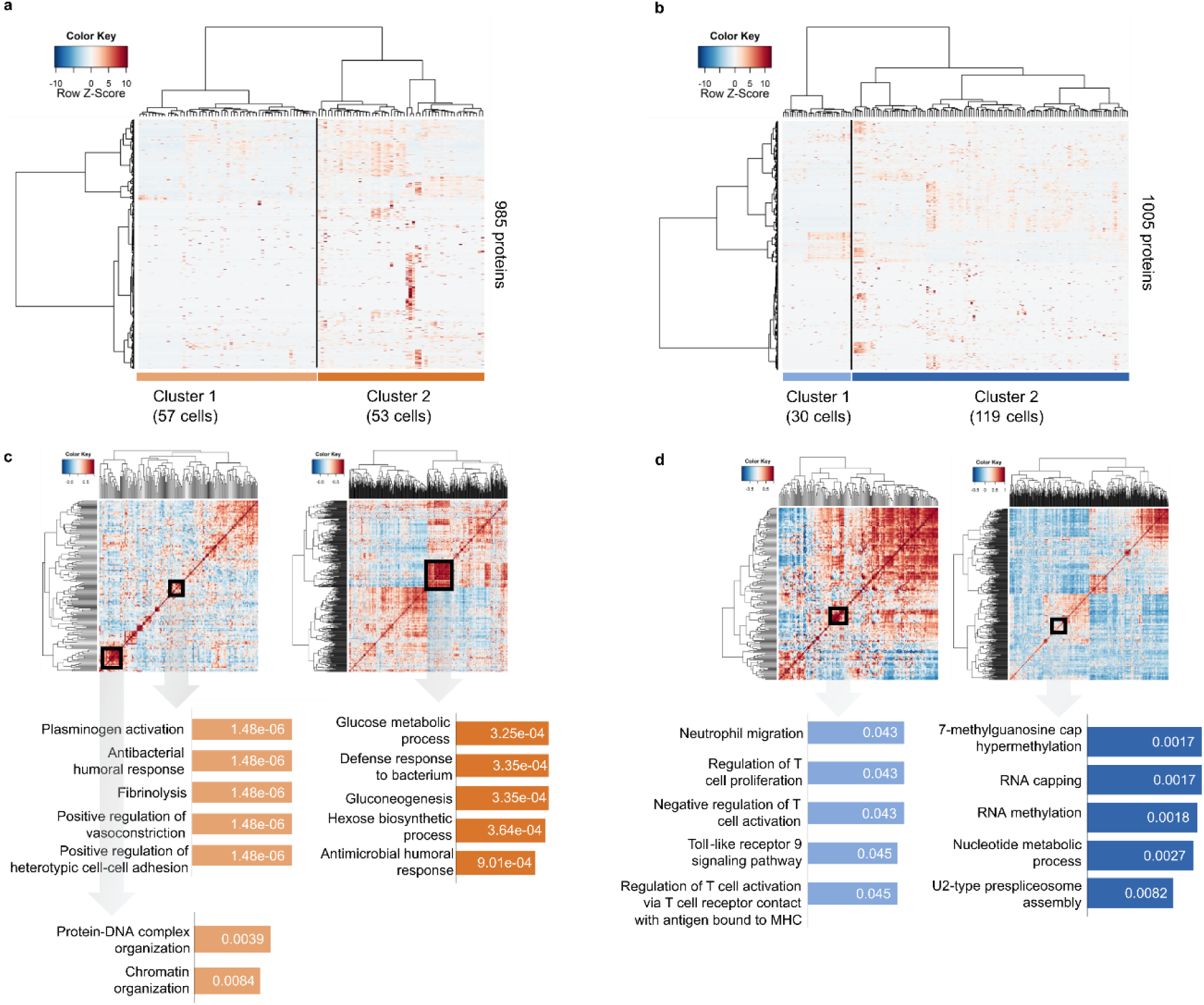
Protein profile and coexpression in saliva and tear single cells. A, B) Heatmap plots were generated with the ‘heatmap.3’ function in the R environment via the Kendall correlation and Ward linkage for the tear fluid dataset (n=149 samples), and the Spearman correlation and Ward linkage for the saliva dataset (n=110 samples). For the heatmaps, datasets were filtered for at least 1 valid value, resulting in 1,005 quantified protein groups in the tear fluid dataset and 985 quantified protein groups in the saliva dataset. C, D) Heatmap of Pearson correlation coefficients (R) derived from pairwise comparisons of the proteins identified in the tear fluid and saliva single-cell data. Protein intensity values were used to calculate the correlation coefficient using the Perseus software v2.1.3.0, and a heatmap was constructed using the R language with the function ‘heatmap.3’. The dendrogram was generated using Euclidean distance with complete linkage. The five most enriched Gene Ontology (GO) biological processes for the highlighted clusters are shown. An adjusted p-value is indicated for each term in the enrichment analysis performed using the Enrichr webtool.

Interestingly, PGs from saliva single cells in Cluster 1 were enriched in biological processes related to vascular integrity, immune defense, and cellular interactions, as indicated by highly significant adjusted p-values (1.48E-06). In contrast, Cluster 2 included PGs associated with metabolic and immune-related functions, indicating roles in energy regulation and microbial defense, with adjusted p-values of approximately 3.25E-04 (**Figure 3c**). Single-cell coexpression analysis of the tear fluid dataset revealed that the PGs in Cluster 1 were predominantly associated with immune-related biological processes, indicating a significant role for these PGs in immune modulation and cellular communication (adjusted p-values: 0.043–0.045) (**Figure 3d**). In contrast, the PGs from Cluster 2 were enriched in processes related to RNA metabolism and assembly, indicating a key role in RNA processing and nucleotide synthesis (adjusted p-values: 0.0017–0.0082). Overall, this analysis allowed us to investigate the data from a distinct perspective and to highlight the most important PGs and their respective biological processes coexpressed in individual cells.

We subsequently analyzed the Gene Ontology (GO) biological process annotations of the PGs in saliva and tear single cells exclusively using experimental annotations retrieved from UniProt. We employed two strategies: (a) count, which involves evaluating the represented GO terms on the basis of their frequency in PGs, and (b) sum, which involves assessing the contribution of the PGs by summing the MS1 intensities (PG1) of the GO terms (illustration of the analysis in **Supplementary Figure 9**).

The saliva and tear single-cell proteomes were associated with 2,052 and 2,059 terms, respectively. To select a representation of cells on the basis of GO terms through UMAP visualization, we evaluated the neighborhood preservation metric Sequence Preservation View^19^ of the UMAP results for 4,000 combinations of initial point placements, distance metrics (cosine and Euclidean), and initialization methods (PCA and random). Finally, we explored the overrepresentation of biological processes by analyzing the frequency of GO terms (counts) (**Figures 4a**, **b**) and the summed intensity of PGs (sum) (**Figures 4c**, **d**) associated with specific GO terms. The UMAP projections indicate that protein functions (GO) can group single cells. Notably, for both saliva and tear samples, the contribution of PG intensities (sum) is relevant to cell clustering (**Figures 4c, d**), indicating that several PGs are involved in the same processes, reinforcing the rationale of cell grouping, especially for tear cells. The top 10 biological processes responsible for the grouping are listed in **Figures 4a-d**. We used the SelectKBest function of scikit-learn to identify the features that best explain the clusters, as shown by the colored cells in the UMAP results (**Figures 4a-d**). These data indicate that the proteomes of saliva and tear single cells are different but have complementary biological functions. In saliva, processes such as phagocytosis, epithelial cell differentiation, T-cell activation and mRNA splicing via the spliceosome dominate the GO count and highlight roles in immune defense, metabolic regulation, and cell maintenance. The sum of protein intensities underscores these findings and reveals functions such as cell cycle regulation, platelet aggregation, oxidant detoxification and mRNA splicing, indicating coordinated protein activity in metabolic and structural processes. In contrast, the proteome of tear single cells is involved in a variety of processes, such as plasminogen activation, cholesterol biosynthesis, positive regulation of cell migration and blood coagulation, indicating vascular, apoptotic and transcriptional regulatory functions. The sum of the intensities in tear cells further underscores antiviral responses, keratinocyte activity, the regulation of MAPK signaling, and desmosome assembly, highlighting roles in tissue integrity, immune defense, and epithelial homeostasis. These results indicate that saliva affects metabolic and immunological interactions, whereas tear cell proteomes exhibit greater diversity, with vascular and epithelial functions contributing to tissue maintenance and defense. The analyses of coexpressed and enriched GO terms are complementary; despite their differences, both approaches emphasize the functional significance of protein clusters. Coexpression analysis revealed novel protein relationships, whereas GO analysis provided a contextual biological interpretation of these processes. Furthermore, both methods overrepresented immune processes in both biofluids, especially in saliva biofluids.

**Figure 4.**
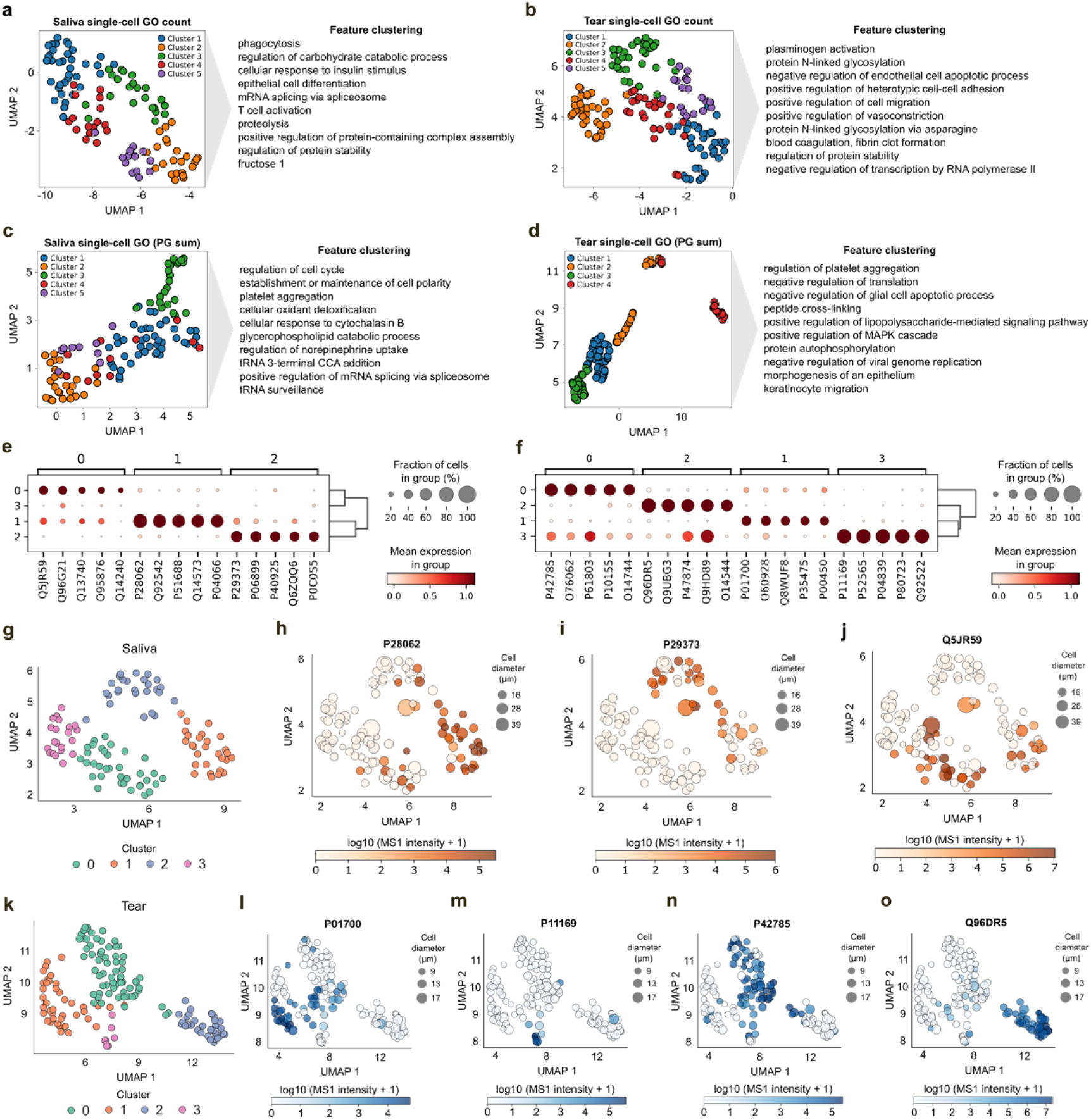
Exploring the biological value of saliva and tear single cells. A, B) UMAP results and annotation list of indicated overrepresented biological processes by GO frequency (counts) and C, D) the sum of the intensity of PGs associated with specific GO annotations. E, F) Scanpy dot plots generated for all markers exported for saliva (E) and tear fluid (F) by each source cluster, with their respective abundances transformed using log10(MS1+1). Normalization was performed utilizing the standard_scale=“var” parameter, ensuring that each protein’s mean abundance was scaled to a range of 0 to 1. G) Leiden clusters in saliva cells, with each cluster represented by a distinct color. H-J) UMAP visualization was employed to represent the Leiden clusters of saliva cells, incorporating marker abundances and cell diameters. Specifically, Q5JR59 was identified as a candidate exclusive to Cluster 0, P28062 to Cluster 1 and P29373 to Cluster 2. No exclusive protein was identified for Cluster 3. K) Leiden clusters in tear cells, with each cluster represented by a distinct color. L-O) UMAP visualization was employed to represent the Leiden clusters of tear cells, incorporating marker abundances and cell diameters. Specifically, P42785 was identified as a candidate exclusive to Cluster 0, P01700 to Cluster 1, Q96DR5 to Cluster 2 and P11169 to Cluster 3.

The single-cell data obtained from the PG and MS1 intensities were also divided into groups of comparable cells using the Leiden algorithm for cell clustering, resulting in clusters on the basis of overall proteome similarity. The individual proteome of each cell was also investigated by analyzing marker genes within clusters, the proportion and mean expression of which were plotted individually for both biofluids (**Figures 4e, f**). The results demonstrated that although a PG may appear exclusive to a community, it is still shared across one or more clusters. It is possible to visualize this behavior in the UMAP color via PG quantification. The Leiden algorithm was used to identify well-connected clusters within the dataset. These four clusters exhibit distinct separations and unique PG abundance profiles, enabling the identification of defining features for each cluster (**Figures 4g, k**).

Despite the diversity of cells, three PGs identified in saliva (**Figures 4h-j**) and four in tear single cells (**Figures 4l-o**) were overrepresented and defined the clusters. The identification of PGs can retrieve potential single-cell types based on clusters of RNA data from The Human Protein Atlas^20–22^. The cluster markers can potentially indicate cell types like neuron cells (high reliability, MTUS2, and Q5JR59), Langerhans cells (medium reliability, PSBM8, and P28062), and connective tissue cells (medium reliability, CRABP2, and P29373). For tear fluid, adipocytes, endothelial cells, monocytes, and neutrophils (both medium reliabilities, PRCP, P42785), plasma cells and B cells (high reliability, IGLV-47, P01700), serous glandular cells (medium reliability, BPIFA2, Q96DR5) and monocytes and neutrophils (medium and low reliability, SLC2A3, P11169) were most represented. Taken together, these results identify relevant proteins that define the clusters and indicate potential cells present in saliva and tear fluid. Notably, processes and cell populations associated with the immune response were overrepresented across the single-cell data analysis as shown in the **Supplementary Figure 10**, with 36 proteins identified in saliva and 38 in tear fluid.

In addition to annotating the functions and potential cell types present in saliva and tear single cells, the corresponding standard gene names of the PGs listed for saliva and tear single cells were compared with the gene names (including isoforms) listed as therapeutic targets associated with FDA-approved drugs (https://www.fda.gov/drugs/science-and-research-drugs/table-pharmacogenomic-biomarkers-drug-labeling; Accessed on February 20^th^, 2025) (**Figure 5a** and **Supplementary Tables 15-17**). Twenty-three therapeutic targets associated with FDA-approved drugs were identified in saliva and/or tear cells (**Figure 5b**). Notably, 30% of these targets are pharmacogenomic therapeutic targets in oncology or metabolic diseases (**Figure 5c**). Among them, LMNA (Prelamin-A/C, UniProt ID P02545) and NPM1 (Nucleophosmin, UniProt ID P06748) were consistently detected in more than 80% of the analyzed cells in both saliva and tear fluid.

**Figure 5.**
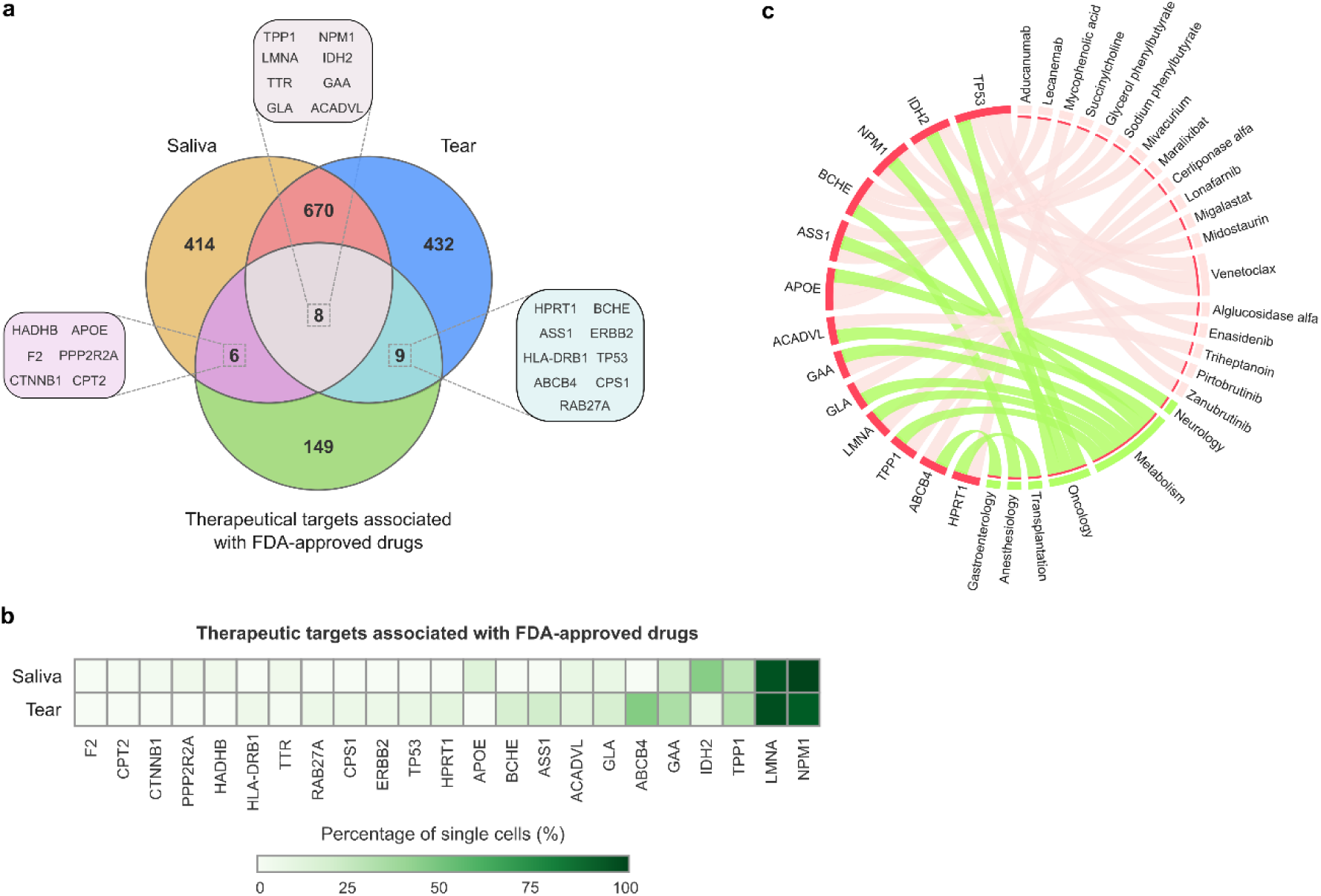
Disease signatures are present in single cells of saliva and tear fluid. A) Venn diagram of the PGs identified in saliva and tear single cells shared with therapeutic targets associated with FDA-approved drugs. B) Heatmap of gene names of the respective FDA-therapeutic targets, on the basis of the percentage (%) presence in cells from biofluids. C) Circle plot of the identified therapeutic targets, FDA-approved drugs, and therapeutic areas. Only 13 biomarkers with ≥9% detection in single cells are included in the plot.

## Discussion

This study demonstrates the feasibility of a label-free SCP approach^6,7,9,23^ for more heterogeneous biological matrices. We explored the effects of single-cell metrics on the biological significance of human saliva and tear cells. To the best of our knowledge, this is the first study addressing these challenges in small-sized cells from fluids, integrating their SCP data with enriched biological processes, clustering algorithms, and therapeutic targets, along with complementary analysis of immunophenotyping using flow cytometry. While this study successfully addressed many challenges in analyzing highly heterogeneous samples, some bottlenecks should be noted. Although pooled samples were used to enhance cell diversity, this approach may have introduced even greater heterogeneity. However, this negative point can help guide the improvement of parameters of cell isolation and search engines, augmenting their ability to handle complex datasets.

One of the main challenges of the SCP workflow is the background or contamination, which becomes particularly relevant when working with low and ultralow inputs^24^. To address this challenge, additional cleaning steps were incorporated into the workflow, both before and after cell isolation, along with different types of blanks throughout the experiment to integrate systematic background contaminants. In our analyses, PGs identified in cells that matched those found in blanks (1-2) were filtered out for the PCA and UMAP results, improving clusterization, in addition to leading to more accurate biological annotations of the cells (**Figures 2h-j** to **5**).

The impact of heterogeneous populations is also reflected in the data analysis. The large global differences between samples can lead to an increase in low-quality peptides within the spectral library. This heterogeneity increases score cutoffs, which in turn hinders accurate protein identification. Thus, after several rounds of data evaluation using Spectronaut, some files were excluded, as represented in **Figures 2c-f**. While removing certain samples may exclude less abundant cell types from analysis, it can increase confidence in identifying well-represented proteomes by narrowing the search space and increasing sensitivity, thus reducing false-negatives due to unnecessary comparisons. This is particularly important when the focus is on the quantification of peptides in different samples^25^. Additionally, the extremely low protein availability of such small cells results in low MS/MS signals, which hampers effective identification via conventional database searches^26^. Furthermore, while the MS2 spectra from the Astral analyzer offer richer data due to its high sensitivity and capacity to detect single ions, this sensitivity can also result in the detection of numerous low-intensity background peaks, including fragmented or scattered ions, which contribute to chemical noise, and thus it also may lead to lower quality identifications^27^.

Another important consideration is that the total number of PGs identified includes all proteins that fulfill the specified filter criteria, although the number of PGs identified and quantified is very similar. Therefore, PGs were not further filtered by the number of cells in which they were detected. In addition, filtering on the basis of protein coverage, number of unique peptides, single hits, and quantification accuracy (e.g., number of points per peak) was not prioritized. Furthermore, the decision to report the data at the MS1 level was mainly driven by the higher number of data points per peak (a median of 6 for saliva and tear samples) at this level, compared to just a median of 2 and 3 at the MS2 level. This choice was also supported by the lower %CVs observed at the MS1 level during standard HeLa digestion. The acquisition method employed ensures longer injection times across broad DIA windows, while enabling comprehensive precursor elution sampling. This method relies on high-resolution MS1 for quantification, a feature also leveraged by Spectronaut. This is an important aspect in the field of SCP, where identification has been prioritized over quantification, although significant efforts continue to be made to improve quantification accuracy. In fact, this happens because Orbitrap scans more slowly and spends more time per scan, capturing more data points per peak, and in parallel, Astral has faster scans and moves quickly between different peptides, thus spending less time on each one, resulting in fewer data points per peak. However, Astral compensates for this issue with its high sensitivity.

Additionally, in this study, the MS acquisition parameters were the same for both biofluid libraries and biofluid single cells. The libraries, however, could have been further optimized, particularly in terms of DIA isolation windows and ITs. Although the high-quality spectral library at the single-cell level has been described to distinguish between normal and cancerous pancreatic cells as previously described^26^, our findings emphasize that the challenges in generating spectral libraries are closely tied to the inherent heterogeneity of cells within biofluids.

To further advance the potential benefits of using cells from noninvasive sources as biofluids, our data were compared with those of FDA-validated therapeutic targets, and twenty-three targets were PGs identified in saliva and/or tear cells (**Figure 5**), reinforcing the potential to greatly increase the applications and contributions of this technique in the treatment of disease. Among the various reported applications, ApoE, identified exclusively in saliva samples, is notable for its use as a target in the treatment of neurological diseases. Among the seven targets associated with metabolic disorders, LMNA has a detection rate of more than 70% in individual saliva and tear cells. NPM1 was similarly detected in both tear and saliva cells, with an average identification rate of over 70%, and it was the only identified target in the present study associated with oncology therapy aside from IDH2. Interestingly, IDH2 was detected more frequently in saliva cells (40%) than in tear cells (9%) and was associated with both oncological and metabolic disorders. While the modulation of protein abundance is not well documented in many cases or is known to be affected by drug treatments, this knowledge can be expanded by using SCP approaches to improve detection methods for potential target proteins and elucidate the interactions between the subtypes of cells that express the therapeutic targets.

Another challenge is the annotation of cell type in SCP, due to the complexity and sparseness of protein quantification data at the single-cell level. Unlike scRNA-seq, for which there are well-established computational tools (e.g. SingleR^28^, Celltypist^29^, scAnnotate^30^, MultiKano^31^), there are no dedicated automated methods for annotation in SCP. Instead, it relies on manual marker-based gating, which is labor-intensive and subjective. The lower feature dimensionality and missing values in mass spectrometry-based SCP further complicate classification^32^. While some scRNA-seq frameworks such as Seurat and Scanpy can be adapted, their effectiveness is limited due to incompatibility with proteomics-specific data normalization. There is a growing need for computational tools tailored to SCP as well as comprehensive protein marker databases^33,34^, especially for underexplored areas such as single-cell data obtained from biofluids. Future advances will likely include the development of specialized algorithms or the adaptation of scRNA-seq methods to improve automated and accurate annotation of cell types in SCP.

Despite significant advancements in SCP through the one-pot workflow integrating the cellenONE X1, Vanquish Neo UHPLC, and Orbitrap Astral instruments, several bottlenecks remain. These include the need for increasing the throughput of cell isolation, particularly in complex microenvironments, and possibly performing presorting via flow cytometry to enrich the target populations, enhancing imaging techniques for morphology-based cell selection, automating the selection process to improve efficiency, and mitigating the presence of contaminants. Additionally, the development of search algorithms is crucial for effectively analyzing diverse cell populations and improving the built spectral libraries. Finally, algorithms for cell type annotation based on protein cluster markers still need to be implemented to increase the reliability of annotation.

Finally, this study demonstrated the robustness of the SCP workflow to reveal cell-specific protein profiles in saliva and tear fluid, providing valuable insights into both biological functions and technical advances. By integrating coexpression analysis, GO term enrichment, protein clustering, and cell annotation, we demonstrated that functional signatures could be effectively retrieved from salivary and tear single-cell datasets. Taken together, these results highlight the ability of SCP to provide both functional and diagnostic insights by linking protein abundance to cell-specific conditions, providing a promising avenue for the advancement of personalized medicine through the study of biofluids and other complex microenvironments.

## Methods

### Preparation of the Pierce™ HeLa Protein Digest Standard for benchmarking

All preparations were conducted using low-retention tips (Maxymum Recovery, Axygen), low-protein-binding microcentrifuge tubes (Thermo Scientific, #3448), and SureSTART Level 3 glass microvials (Thermo Scientific, #6PSV9-V1T). The HeLa protein digest (Pierce HeLa Protein Digest Standard, Thermo Scientific, #88329) was resuspended in a solution containing 0.1% formic acid (FA) and 0.015% n-dodecyl β-D-maltoside (DDM) (Sigma, #D4641) and then sonicated at 40 Hz for 5 min using an FS20 sonicator (Fisher Scientific). The final protein concentration was adjusted to 5 ng/µL. Prior to injection into the LC‒ MS/MS system, glass vials were rinsed with the corresponding peptide solutions. The injection volumes ranged from 50 to 200 nL.

For the commissioning of instruments, sample transfer was minimized to avoid contamination or losses, and all plastics were low-binding to preserve sample integrity, as previously indicated^35^. All the samples were freshly prepared, and the instruments were thoroughly cleaned and calibrated prior to use. Analytical columns were conditioned by tryptic protein digestion via sequential injections of 10–100x the desired input or following the manufacturer’s instructions.

### Preparation of HeLa cells for benchmarking

HeLa cells (CCL-2) were purchased from the Rio de Janeiro Cell Collection (Banco de Células do Rio de Janeiro -BCRJ, Brazil) and were authenticated by the ATCC and tested for negative mycoplasma contamination. The cells were maintained in Dulbecco’s modified Eagle’s medium (DMEM) supplemented with 10% fetal bovine serum (FBS) and 1% antibiotics (penicillin‒streptomycin) at 37 °C and 5% CO_2_. For SCP analysis, the cells were cultured in a medium flask (T75) and collected when they reached 80–90% confluence. Next, the cells were washed twice with 1X phosphate-buffered saline (PBS), trypsinized and incubated with the Zombie NIR™ Fixable Viability Kit at a final concentration of 1:250 for 15 min at room temperature, then washed twice with 1X PBS and resuspended in previously filtered (Millipore® Steritop® Vacuum Bottle Top Filter, Merck, #SCGPS02RE) and degassed 1X PBS.

### Ethical approvals and biofluids collection

This study included biofluid samples from healthy individuals (pool of n=10) and was approved by the Ethics Committee of the Piracicaba Dental School (FOP), São Paulo, SP, Brazil, through protocol CAAE: 54995522.0.0000.5418. All participants provided written informed consent. The study was conducted following the Declaration of Helsinki and was performed following the Strengthening the Reporting of Observational Studies in Epidemiology (STROBE) statement^36^. The procedures used for saliva and tear fluid sampling and annotation were performed following the guidelines and experimental protocols approved by ethics committees. The information collected from the individuals included the following: sex (n= 7 females, n=3 males) and mean age (n=4 >38.7, n=6 < 38.7).

Saliva samples were collected from individuals advised not to eat or brush their teeth for at least 1 h before sample collection. First, the mouths of the 10 volunteers were rinsed with 5 mL of drinking water, and then approximately 2 mL of unstimulated saliva was collected in a 15 mL sterile plastic tube, with collection lasting up to 5 min^37^. To obtain the saliva pool (n=10), approximately 2 mL of saliva from each volunteer was combined in a 50 mL sterile plastic tube, and the saliva was harvested by centrifugation at 20,800 × g for 10 min at 4 °C to recover the cellular content (modified from Winck et al.^37^, Busso-Lopes et al.^13^, and Jiao et al.^38^). Afterward, the supernatant with clear fresh saliva was collected and transferred to a new tube, and both the supernatant and the pellet were maintained on ice until Ficoll cell extraction.

To collect tear fluid samples, 80 µL of sterilized fresh saline solution was added to each eye with a 200 µL micropipette, and the excess saline solution was collected with a capillary tube placed on the lateral corner of the eye. The tear fluid was recovered from the capillary tubes by blowing air in the tube with a 200 µL micropipette, and the solution from each eye was transferred inside the same 1.5 mL microtube. Saliva and tear fluid were maintained on ice until further use. To collect tear fluid samples, 80 µL of sterilized fresh saline solution was added to each eye with a 200 µL micropipette, and the excess saline solution was collected with a capillary tube placed on the lateral corner of the eye. The tear fluid was recovered from the capillary tubes by blowing air in the tube with a 200 µL micropipette, and the solution from each eye was transferred inside the same 1.5 mL microtube. Saliva and tear fluid were maintained on ice until further use.

### Cellular extraction from saliva and tear fluid

Saliva mononuclear cell content was obtained by Ficoll extraction as previously described (Busso-Lopes et al.^13^) with modifications. Saliva cell pellets recovered by centrifugation were resuspended in 4 mL of PBS and diluted in DMEM without FBS (1:1). Next, 4 mL of Ficoll 1077 (Ficoll-Paque® PLUS Medium) was added, and the mixture was centrifuged at 900 × g for 30 min. To ensure cell recovery, we also diluted 4 mL of the supernatant of saliva in DMEM (from previous 20,800 × g centrifugation) and performed Ficoll extraction in parallel. The cellular ring was removed, placed in a clean tube and washed three times with 1X PBS at 400 × g for 10 min. Final saliva cells were obtained via a combination of both Ficoll recoveries. Tear fluid cells were obtained by centrifuging approximately 500 µL of tear fluid in 1X PBS in 1.5 mL RNase-DNAse-free sterilized tubes, as described, at 400 × g for 5 min at room temperature. Saliva and tear fluid cells were subsequently incubated with the Zombie NIR™ Fixable Viability Kit at a final concentration of 1:250 for 15 min at room temperature. This was followed by a single wash with degassed and filtered 1X PBS prior to analysis by both cellenONE X1 and flow cytometry.

### Detection and isolation of cells assisted by cellenONE X1

The cellenONE® X1 robot (Cellenion®, Lyon, France) was employed for the automatic dispensing of reagents and for cell isolation on the basis of morphometric and fluorescence parameters. All the solutions were prepared using MS-grade reagents, degassed prior to use and loaded into a medium-sized uncoated piezoelectric dispensing capillary (PDC). A drop check was performed every 12 wells to ensure dispensing accuracy. The master mix, consisting of 100 mM TEAB, 0.05% DDM (Sigma, #D4641), 1% dimethylsulfoxide (DMSO, Sigma, #472301), 0.01% trypsin enhancer (ProteaseMAX, #V2072), and 3 ng/µL trypsin (RapiZyme, #186010108), was dispensed into each well of a clean Eppendorf LoBind 384-well PCR plate (Merck, #EP0030627300) in two consecutive rounds, each dispensing 500 nL. Following mapping and analysis of the cells, cell isolation was performed on the basis of parameters set on the cellenONE X1. The transmission channel was initially used for cell selection, followed by fluorescence channel analysis using a red LED for negative selection of viable cells (T>F). To ensure the detection of all the cell particles, the detection parameters were set to a wide range: 5 to 100 µm in diameter and up to 4 in elongation. The following cell isolation parameters were optimized for each cell population: HeLa cells (n=20) (benchmark) were set to a diameter between 20 and 40 µm with an elongation of up to 1.8, saliva cells (n=200) were set between 5 and 45 µm in diameter with an elongation of up to 1.95, and tear cells (n=200) ranged from 5 to 20 µm in diameter with elongation of up to 1.95. During the two runs (master mix dispensing and cell isolation), the 384-well plate was maintained at 10 °C and 45% humidity. Following, the plate was incubated at 50 °C for 2 h. The humidity was set to 85% to minimize evaporation, and hydration steps were applied with 500 nL of water. The second round of hydration was replaced with 500 nL of 3 ng/µL trypsin resuspended in water, as described by Matzinger et al.^23^ and Bubis et al.^6^. After 2 h of incubation, 4 µL of 0.1% FA was added to each well, using an electronic pipette repeater, to quench the trypsin activity. The plate was then dried in a Thermo SC250EXP SpeedVac concentrator at room temperature and stored at -20 °C until injection. Wells containing multiple cells were processed through the same workflow to construct sample size-comparable spectral libraries^39^. For the HeLa benchmarks, ten isolated cells were processed in two replicates, whereas for saliva and tear, eight wells per condition were preprocessed, containing 6 and 20 cells, each. The two replicates with the greatest number of identifications were selected for library construction. As experimental controls for SCP, during cell isolation, we included blank wells on the cell isolation plates (n=6 for HeLa benchmarks; n=12 for saliva and tear fluid), which contained only the master mix without any isolated cells. A plate with only blank wells was also incubated under the same conditions as the cell-containing plates on the next day of the experiments with isolated cells (n=10). All the SCP experiments were performed after cleaning the equipment with 70% ethanol (“SOP on a 14-day” task on the cellenONE X1).

For protocol optimization with HeLa cells, we included an additional cleaning step of PDC called “Sterilization”, before and after cells isolation, with three solutions used in sequential (0.5% sodium hypochlorite, 3% hydrogen peroxide and 70% ethanol). For biofluids, we employed a new medium sized PDC to avoid any cross contamination with remaining HeLa cells and the PDC was cleaned with 70% ethanol before and after cells isolation. Also, the ultrapure water being used in the cellenONE X1 system bottle was replaced with MS grade water.

### LC‒MS/MS

The sample wells were first randomized using the *base package* (version 2.4.6.26) in the R environment (version 4.3.2) across four acquisition batches to replenish chromatographic solvents and perform sampler cleaning. Next, the samples in the 384-well plates were resuspended in 4.0 µL of 0.1% AF, and 3.5 µL was analyzed using a Vanquish Neo UHPLC chromatographic system coupled with an Orbitrap Astral mass spectrometer (Thermo FisherScientific, Germany). Chromatographic separation of peptides was performed on a 25 cm × 75 µm ID, 1.7 µm C18 Aurora Ultimate TS column, with all separations carried out using a single-column configuration. The column oven (IonOpticks) was set to 50 °C. For single-cell-derived peptides, the autosampler and injection valves were configured for direct injection, with fast loading enabled. The system was set to a maximum pressure of 1,200 bar and a loading volume of 1 µL, and a 15 µL injection loop was used to sample from a 384-well plate.

Single-cell-derived peptides were acquired in positive mode using the FAIMS Pro interface, with a compensation voltage set to -48 V and a spray voltage of 1,800 V. Orbitrap MS1 spectra were acquired at a resolution of 240,000, with a scan range of 400 to 800 m/z, a normalized automatic gain control (AGC) target of 500%, and a maximum injection time of 100 ms. Data-independent acquisition (DIA) of MS2 spectra was performed on the Astral using the same scan range, with loop control set to 0.6 seconds per cycle, employing varying isolation window widths and injection times. The isolation window width and injection times ranged from 20 m/z and 60–80 ms for single-cell samples to 10 m/z and 20 ms for the HeLa digest concentration curve (**Supplementary Table 14**). Precursor ion fragmentation was achieved using higher-energy collisional dissociation (HCD) with a normalized collision energy (NCE) of 27%. Gradient elution was conducted at a throughput of 50 samples per day (SPD) in an active gradient. The percentage of buffer B (80% ACN in water with 0.1% FA) initially increased from 0 to 40% over 19.5 min at a nominal flow rate of 450-200 nL/min. This was followed by a gradual increase to 99% buffer B from 19.5 to 22 min at a flow rate of 300 nL/min, which was maintained for an additional 5 min. Data acquisition was performed for 24 min from the start of elution. Following data collection, the FAIMS voltage was set to 0 V during the washing and equilibration steps to minimize contamination of the mass spectrometer. Blank runs were performed in the LC‒MS/MS analysis after every 10 samples to minimize accumulation of DDM in the analytical column.

### Data processing and analysis

The raw files were converted to .htrms files via the HTRMS Converter (Biognosys, version 19.2) and processed using Spectronaut (Biognosys, version 19.2.240905.62635) library-free search (directDIA+) with the BGS factory settings. Trypsin/P was selected as the protease, with up to two missed cleavages allowed. Methionine oxidation and N-terminal acetylation were set as variable modifications. If not otherwise stated, the following identification thresholds were applied: precursor Qvalue < 0.01, precursor PEP < 0.2, protein (experiment) Qvalue < 0.01, protein (run) Qvalue < 0.05, and protein PEP < 0.75. Protein identification was carried out against the UniProt human reference proteome (UP000005640, 20,654 entries, downloaded on August 1st, 2024), supplemented with the common Repository of Adventitious Proteins (cRAP – 116 entries) to eliminate common mass spectrometry contaminants, which were filtered out after analysis.

For method evaluation, all samples were analyzed together in a single project with the Pulsar DIA Method Evaluation feature enabled. For benchmarking with directDIA+, samples were treated as replicates of the same condition, employing cross-run normalization and matching between runs. When files categorized as “all files” were processed, all the raw files were included in a single project but were configured under different conditions. Bulk sample replicates based on single-cell isolation were utilized to generate either Pulsar spectral libraries or sample replicates using directDIA+ (matching enhancer or DIA-ME). Blank samples (blanks 1-3) were independently analyzed by directDIA+ by each condition as sample replicates.

Protein intensities were quantified using the built-in MaxLFQ method and normalized via local cross-run normalization. Alternatively, the results from the directDIA+ search were used to construct a communal library, followed by a second round of database searching using the default parameters. Ratio-based normalization of summarized precursor intensities was then performed using directLFQ for biofluid comparisons such as for PCA^40^.

### Immunophenotyping

The cells were washed with 300 µL of filtered 1X PBS by centrifugation at 400 × g for 5 min and incubated for 10 min with 5% FC blocking solution, followed by 10 min of incubation with the antibodies of interest (diluted in filtered 1X PBS at the volume established in the manufacturer’s instructions or determined via an antibody titration assay). The antibodies used for the staining of saliva mononuclear or tear cells were anti-CD45 (Biolegend; USA; clone HI30; FITC-conjugated; 1:100 dilution), anti-CD3 (Biolegend; USA; clone UCHT1; PE-conjugated; 1:100 dilution), anti-CD4 (BD Horizon; USA; clone: L200; PerCPCy5.5-conjugated), 1:20 dilution, anti-CD8 (Biolegend; USA; clone: SK1; APC-conjugated; 1:100 dilution), anti-CD19 (BD Horizon; USA; clone HIB19; PE-Cy7-conjugated; 1:100 dilution) and anti-CD19 (BD Horizon; USA; clone: HIB19; FITC-conjugated). The cells were resuspended in 200 µL of filtered 1X PBS for acquisition via flow cytometry (BD FACS Canto II) and analysis via FlowJo^TM^ 10 software (BD Biosciences, USA). At least 10,000 gated events were acquired, ensuring the reliability of the positive populations. Single cells were gated on forward scatter-height (FSC-H) vs. forward scatter-area (FSC-A) and side scatter-height (SSC-H) vs. side scatter-area (SSC-A) plots. The cells were then gated on a plot of viable cells (Zombie NIR). For the analysis of tear cells, viable cells were gated on CD45^+^ cells, followed by CD3^-^ and CD3^+^ cells. CD3^+^ cells were gated on CD4^+^ and CD8^+^ cells, and CD3^-^ cells were gated on CD19^+^ cells. For saliva analysis, the viable cells were gated directly on CD3^-^ and CD3^+^ cells, followed by CD4^+^ and CD8^+^ cells from CD3^+^ cells and CD19^+^ cells from CD3^-^ cells. The population of T and B cells was evaluated by the percentage of the cells.

### Post-analysis and statistics

To evaluate the batch effect, PCA calculations were performed using the scikit-learn v1.3.0 PCA implementation, and for the plots, the packages Seaborn v0.13.2 and Matplotlib v3.8.0 were employed using Python v3.9.18. A least squares linear regression model was used to assess the relationship between the cell diameter and the number of PGs identified per cell. This strategy reduces the residual sum of squares between the observed targets in the dataset and the targets predicted by the linear approximation. The *LinearRegression* method from scikit-learn was used for this analysis.For data visualization, heatmaps with z score values of protein intensities were generated using the on-line available R function ‘heatmap.3’ (https://github.com/obigriffith/biostar-tutorials/tree/master/Heatmaps). To evaluate coexpressed protein clusters, we used protein intensity values to calculate the correlation coefficient using the Perseus software v2.1.3.0^41^ and constructed a heatmap using the R language with the function ‘heatmap.3’. The dendrogram was generated using Euclidean distance with complete linkage. GO enrichment analysis was performed using the Enrichr^42^ with adjusted p-value < 0.05 chosen as the significance threshold. The entire human proteome GO annotation file was used as a reference set. A heatmap for visualization of FDA biomarkers in cells from saliva and tear fluid was generated using Python’s Seaborn Library (function “heatmap”).

An in-house program written in Perl (5.36.0) was used to download data from UniProt and extract GO terms related to each PG of saliva and tear cells. UMAP projections were performed with in-house programs written in Python (3.11.2) using the package UMAP (0.1.1)^43^ clustering using the package LeidenAlg^44^ (0.10.2) and feature importance using the package scikit-learn (1.6.1). Neighborhood preservation for projections was evaluated using an in-house program written in Python. The data were divided into groups of comparable cells using the Leiden algorithm^44^ resulting in clusters on the basis of overall proteome similarity. Leiden is included in the Scanpy toolkit^45^, and version 1.10.3 was used in our analysis with all the default parameters, varying only the resolution value. To visualize the different clustering results obtained, a UMAP was generated using the Python package umap-learn v0.5.3^43^. Thereafter, the markers were analyzed to determine their associations with known biological features, either as specific types or as cell states, such as comparisons between activated and quiescent states. Next, the protein accession numbers were searched against the Human Protein Atlas^20–22^, on the basis of the RNA expression profiles of single cells and deconvolution of bulk transcriptomics data, the annotated cells were retrieved with their related confidence. Enrichment of the list of PGs from the HeLa, saliva and tear cell proteomes was performed using the Enrichr^42^ with adjusted p-value < 0.05 chosen as the significance threshold. The entire human proteome GO annotation file was used as a reference set. The Blood Atlas, which is part of the Human Protein Atlas^20–22^, was used to annotate the cellular origin of the identified proteins. Sets of enriched cell types were extracted from the Blood Atlas, comprising a total of 727 genes in B cell, 1,395 genes in T cell, 472 genes in NK cells, 962 genes in monocyte, 1990 genes in granulocyte, and 1,139 genes in dendritic cell datasets. Circle plot imaging was performed using Circos online software^46^.

## Data availability

The mass spectrometry proteomics data, along with the raw data from the single-cell isolation experiments, have been deposited in the ProteomeXchange Consortium (http://proteomecentral.proteomexchange.org) via the PRIDE partner repository under the dataset identifier PXD060929.

## Acknowledgments

This work was supported by FAPESP grants EMU 22/11476-5 and 2018/18496−6 to AFPL, 2018/15535-0 to DCG and scholarships 23/12076-3 to HMAP, 23/16823-8 to JGM, 22/12815-8 to DF, 23/16780-7 to EC, 23/16779-9 to Fabio P, 22/14348-8 to GC, and CNPq grants 310392/2021−7 to AFPL. This work was also partially supported or had resources from the Brazilian Federal Government provided to the Brazilian Center for Research in Energy and Materials (CNPEM), a private nonprofit organization under the supervision of the Brazilian Ministry for Science, Technology, and Innovation (MCTI). This study was conducted at the Mass Spectrometry Laboratory of the Brazilian Biosciences National Laboratory (LNBio), which is part of CNPEM, during the commissioning of instruments and methods. We extend our thanks to Dr. Fernanda Salvato from Thermo Scientific for fruitful discussions on improving the LC‒MS/MS methods and Erison Santos from Faculdade de Odontologia de Piracicaba, UNICAMP, for helping optimize tear sample collection.

## Author contributions

JGM, HMAP and AFPL conceptualized the study. MAL and ASS collected the saliva and tear samples. JGM and HMAP conducted the experiments. JGM, HMAP, and AFPL contributed to optimizing and performing the single-cell analysis using cellenONE X1, Vanquish Neo UPLC, and Orbitrap Astral. JGM carried out proteomics searches. Post-search analysis was performed by JGM, HMAP, FMP, CMC, DCG, GC, GT, and AFPL. DLF and EC conducted flow cytometry experiments. AFPL conceived and supervised the study and acquired funding. All authors reviewed, edited, and approved the final version of the manuscript.

## Additional information

No new unique reagents were generated in this study.

## Abbreviations and nomenclature

SCP: single-cell proteomics; LC‒MS/MS: liquid chromatography‒tandem mass spectrometry; FDR: false discovery rate; DIA: data-independent analysis; PDC: Piezo Dispense Capillary; ME: matching enhancer; DIA: data-independent analysis; CV: coefficient of variation; GO: gene ontology; UMAP: Uniform Manifold Approximation and Projection; PCA: Principal Component Analysis; scRNA-seq: Single-cell RNA sequencing.

## Lead Contact

Further information and requests for resources and reagents may be directed to and will be fulfilled by Adriana Franco Paes Leme.

