## supplementary figures for "Single-cell proteomics workflow for characterizing heterogeneous cell populations in saliva and tear fluid"

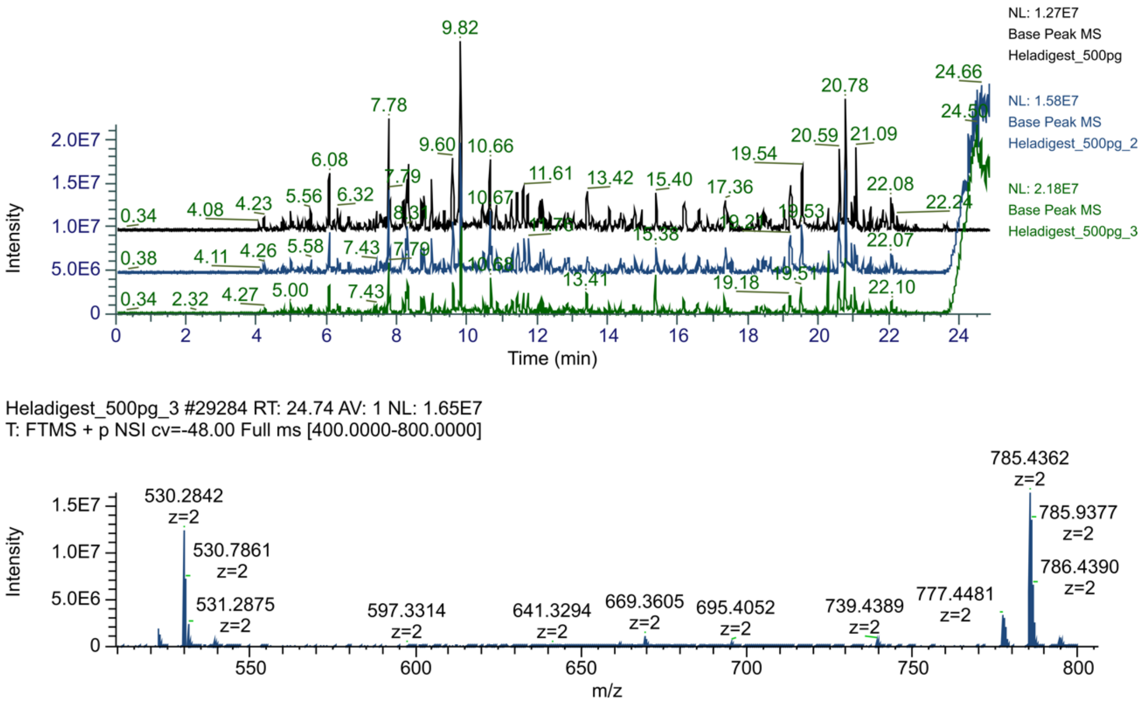


**Supplementary Figure 1. Late elution of surfactant.** Upper panel shows three repeated injections of 500 pg (1 µg/µL) HeLa digest in 0.1% formic acid (FA) and 0.015% n-dodecyl-β-D-maltoside (DDM) demonstrated the selective elution of the surfactant. The overlaid base peak chromatograms of the first (black), second (blue), and third (green) replicate injections reveal the progressive accumulation of charged species at m/z 530.28 and 785.43 (lower panel spectra) attributed to DDM reagent, emphasizing its late elution profile**.**

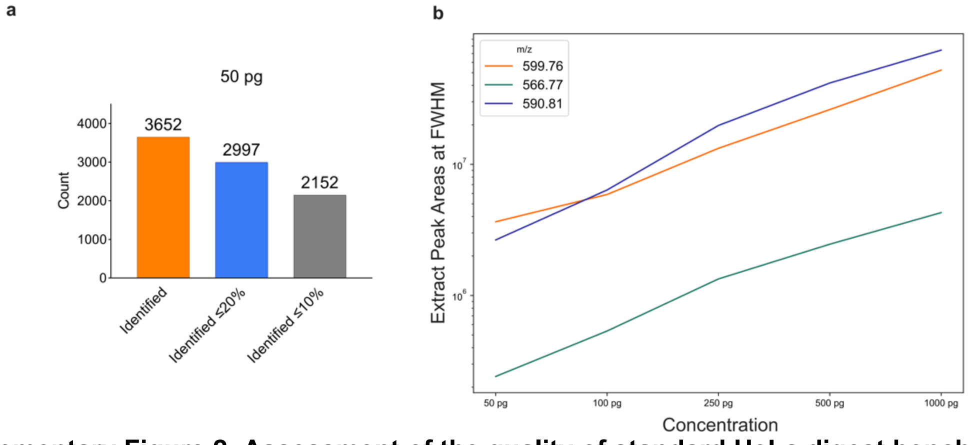


**Supplementary Figure 2. Assessment of the quality of standard HeLa digest benchmarks.** A) Number of identified protein groups in 50 pg of HeLa digest with the percentage of coefficients of variation (%CV) below the specified thresholds (directDIA+, three replicates)**.** B) Measurement of the extracted area at full width at half maximum (FWHM) for three diagnostic peptides across varying injection concentrations.

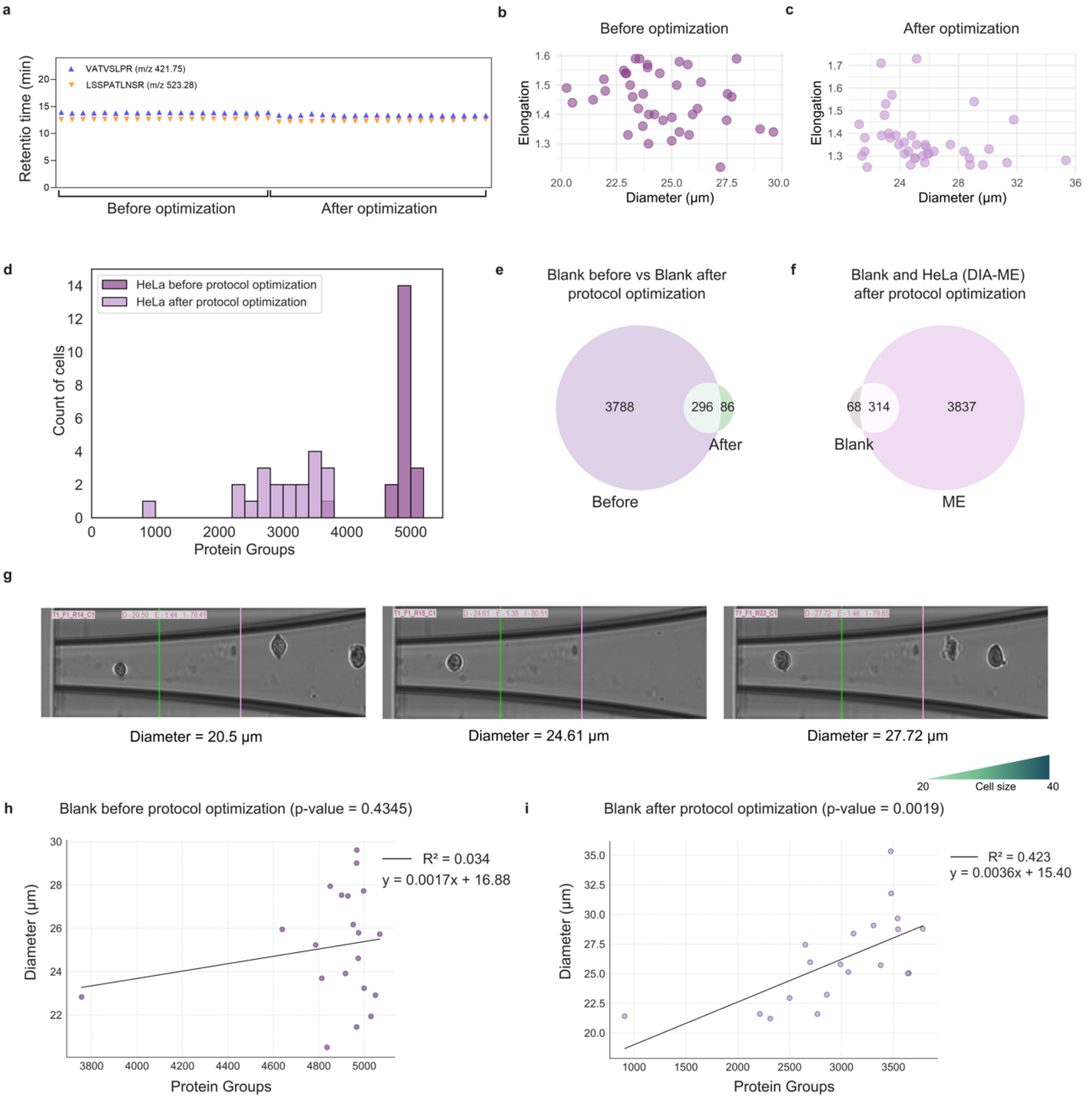


**Supplementary Figure 3. Trypsin efficiency, morphometric parameter analysis and blank controls of HeLa cells before and after cleaning protocol optimization.** A) Evaluation of the reproducibility of two selected autolytic tryptic peptides in individual single HeLa cell runs (n=20), before and after cleaning protocol optimization. B, C) Diameter and elongation information of isolated HeLa cells before and after cleaning protocol optimization. D) Histograms of PGs identified in HeLa cells before and after cleaning protocol optimization and count of cells (cRAP PGs were removed from this analysis). E) Intersection of PGs identified in the blanks before and after protocol optimization. F) Intersection of PGs identified in blanks and single HeLa cells with directDIA+ matching enhancer (ME) after protocol optimization. G) Representative images of isolated cells on the basis of the range of diameter chosen for isolation. H) Linear regression of HeLa cells before cleaning protocol optimization, showing no significant p-value (>0.05) and a low $R^2$ value (0.034) (cRAP PGs were removed from this analysis). I) Linear regression of HeLa cells after protocol optimization, showing a significant p-value (<0.05) and a higher $R^2$ value (0.42) (cRAP PGs were removed from this analysis).

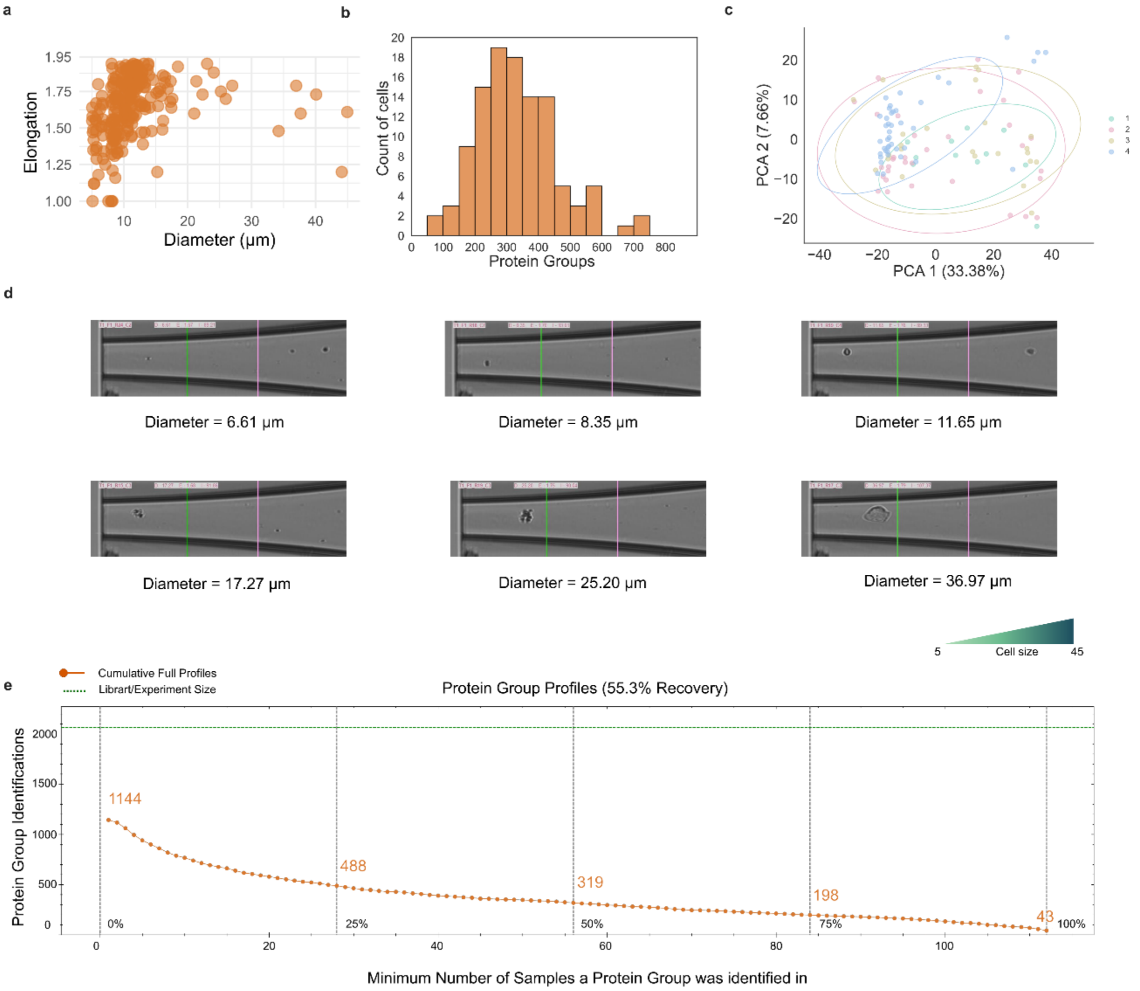


**Supplementary Figure 4. Morphometrics of saliva single cells and PG identification.** A) Diameter and elongation information of isolated cells from saliva. B) Histogram of PGs identified and count of cells (cRAP PGs were removed from this analysis). C) Principal component analysis (PCA) evaluating the batch effect through LC–MS/MS runs (4 batches), with each dot representing a cell. D) Representative images of cells according to the range of diameter chosen for isolation. E) Data completeness at the PG level for all the saliva single-cell files after filtering the raw files (graphic generated by Spectronaut software; cRAP PGs were not removed from this analysis).

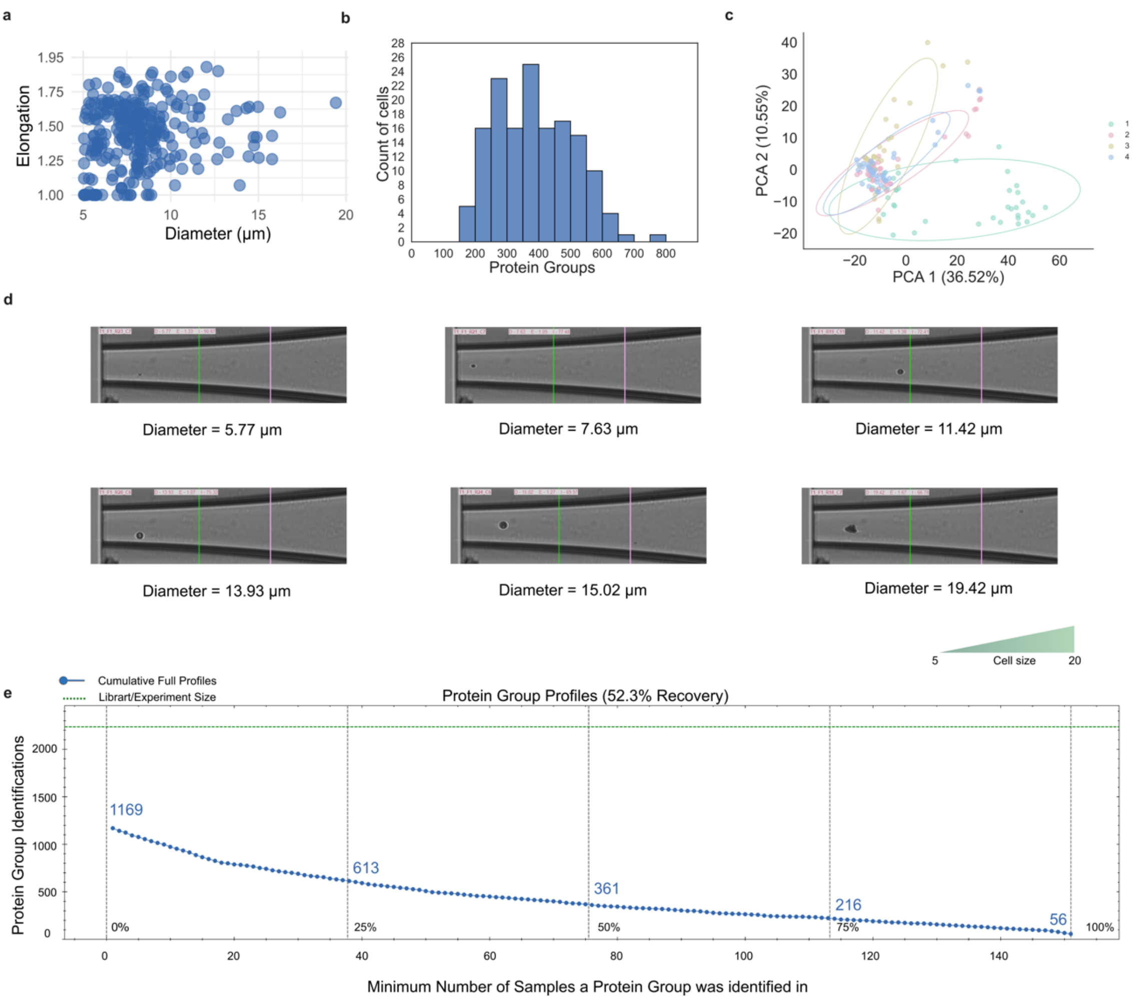


**Supplementary Figure 5. Morphometrics of tear single cells and PG identification.** A) Diameter and elongation information of isolated cells from tear. B) Histogram of PGs identified and count of cells (cRAP PGs were removed from this analysis). C) Principal component analysis (PCA) evaluating the batch effect through LC–MS/MS runs (4 batches), with each dot representing a cell. D) Representative images of cells according to the range of diameter chosen for isolation. E) Data completeness (n=149) at the PG level for all the tear single-cell files, filtering raw files (graphic generated by Spectronaut software; cRAP PGs were not removed from this analysis).

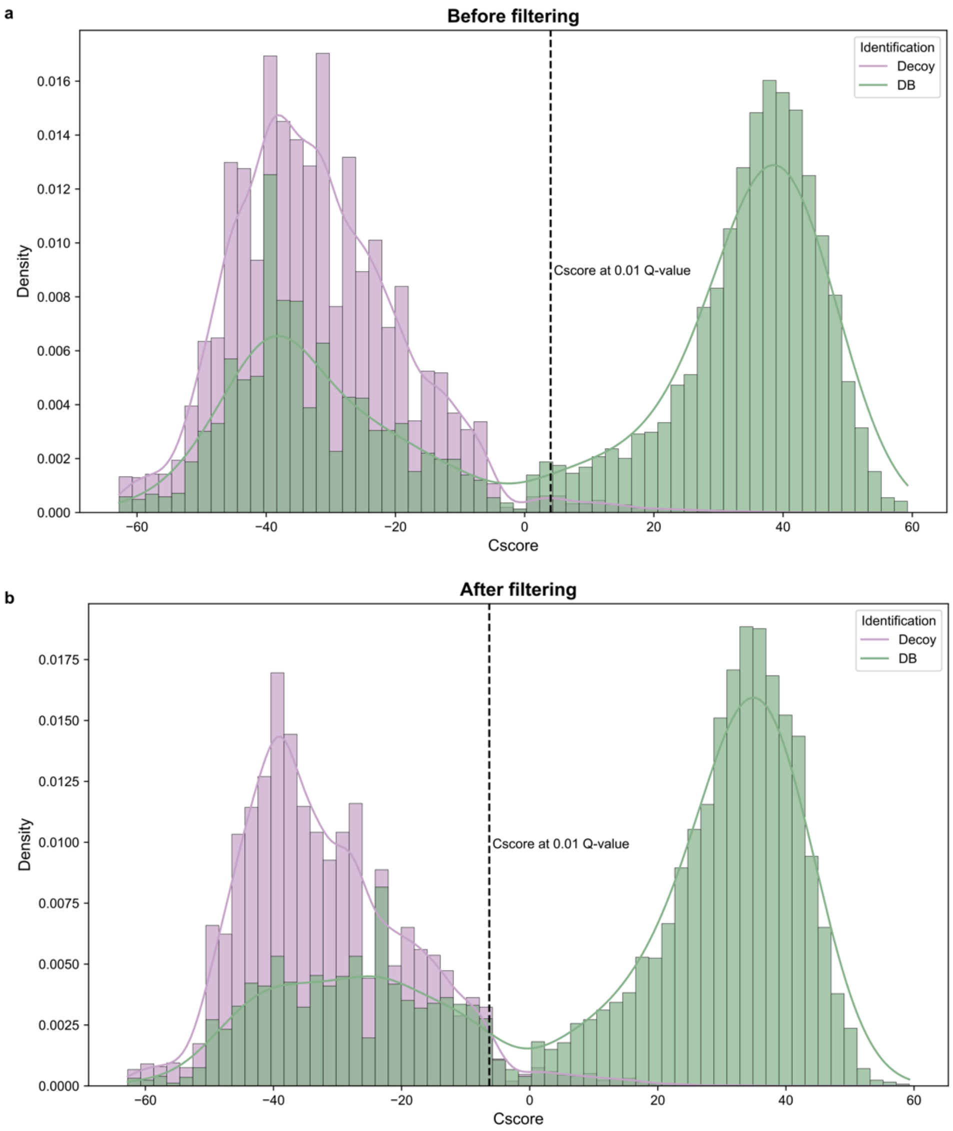


**Supplementary Figure 6. Representative histograms of identifications in a single saliva sample.** Histograms showing the number of identifications in the database and decoy, A) before and B) after optimizing on-the-fly spectral library searches. The data highlights a significant improvement in true positive hits following optimization.

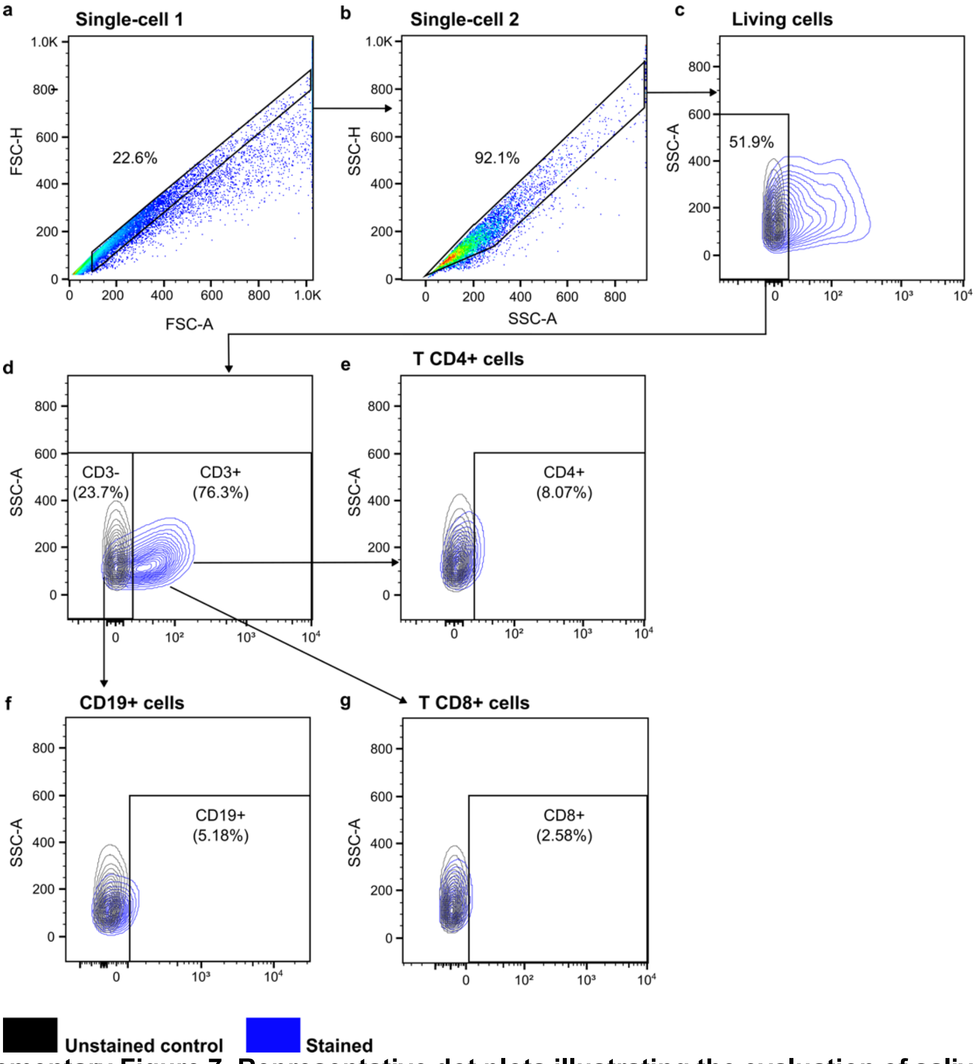


**Supplementary Figure 7. Representative dot plots illustrating the evaluation of saliva cells from 10 healthy individuals obtained through Ficoll extraction**. A) Identification of singlets using FSC-H × FSC-A. B) Verification of singlets using SSC-H × SSC-A. C) Identification of viable cells. D) Discrimination between CD3- and CD3+ T cells. E) Detection of CD4+ T cells gated on CD3+ T cells. F) Identification of CD19+ cells gated on CD3- T cells. G) Detection of CD8+ T cells gated on CD3+ T cells**.**

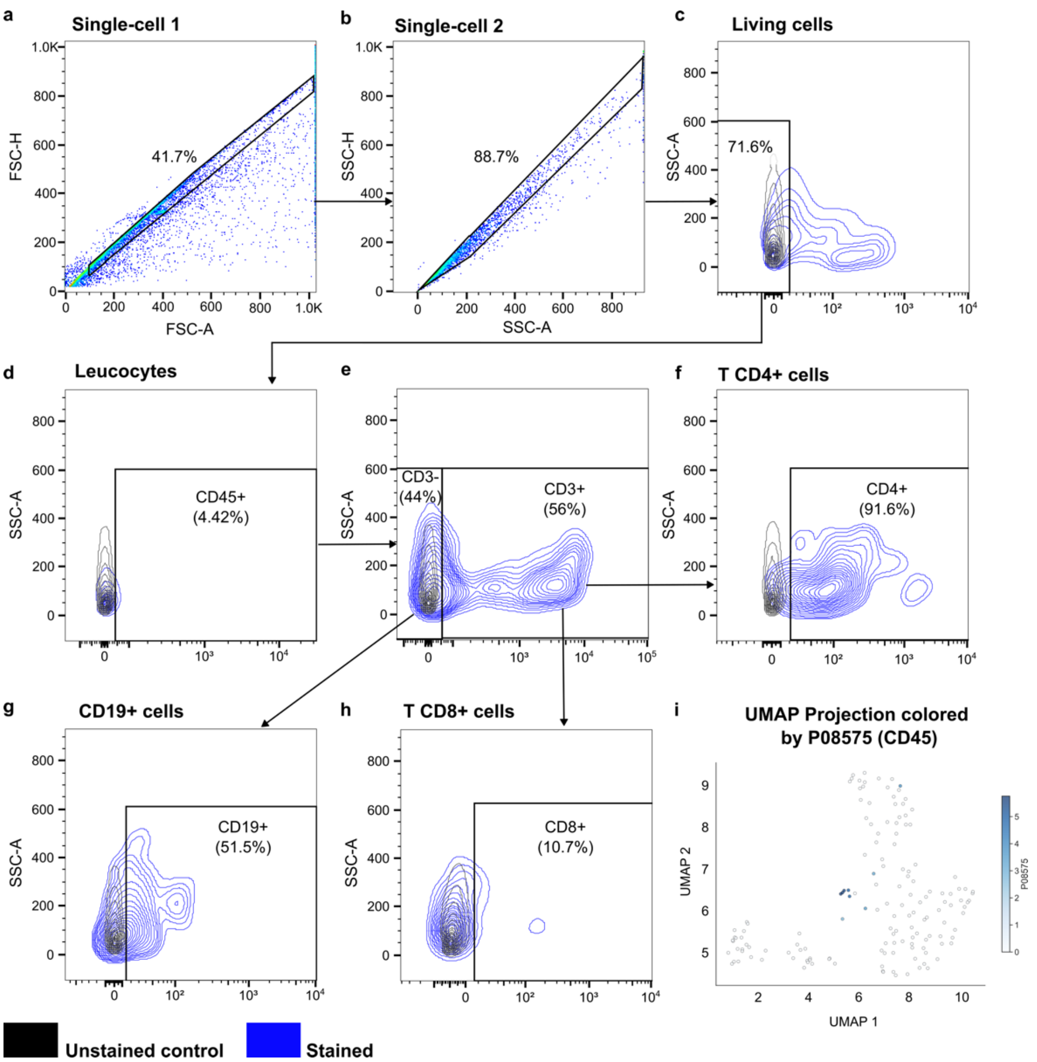


**Supplementary Figure 8. Representative dot plots for the evaluation of tear cells from 10 healthy individuals.** A) Identification of singlets using FSC-H × FSC-A. B) Verification of singlets using SSC-H × SSC-A. C) Identification of viable cells. D) Detection of CD45+ leukocytes. E) Discrimination between CD3- and CD3+ cells. F) Detection of CD4+ T cells gated on CD3+ cells. G) Detection of CD19+ cells gated on CD3- T cells. H) Detection of CD8+ T cells gated on CD3+ cells. I) UMAP visualization of the protein CD45 (P08575) identified in tear single cells, with each dot representing a single cell. The cells were colored according to protein quantitation (log10+1 (LFQ intensity)).

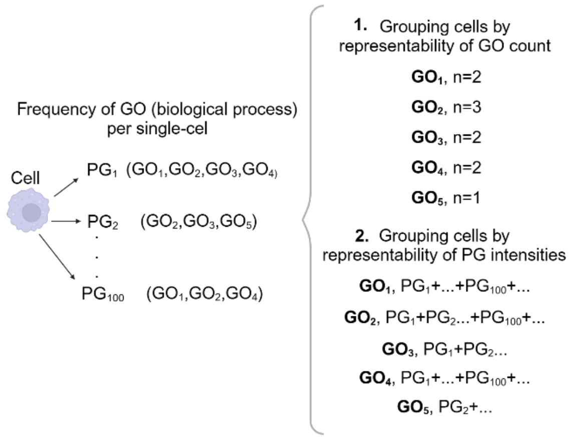


**Supplementary Figure 9. Illustration of the strategies used to analyze the GO biological process of PGs in saliva and tear single cells.**

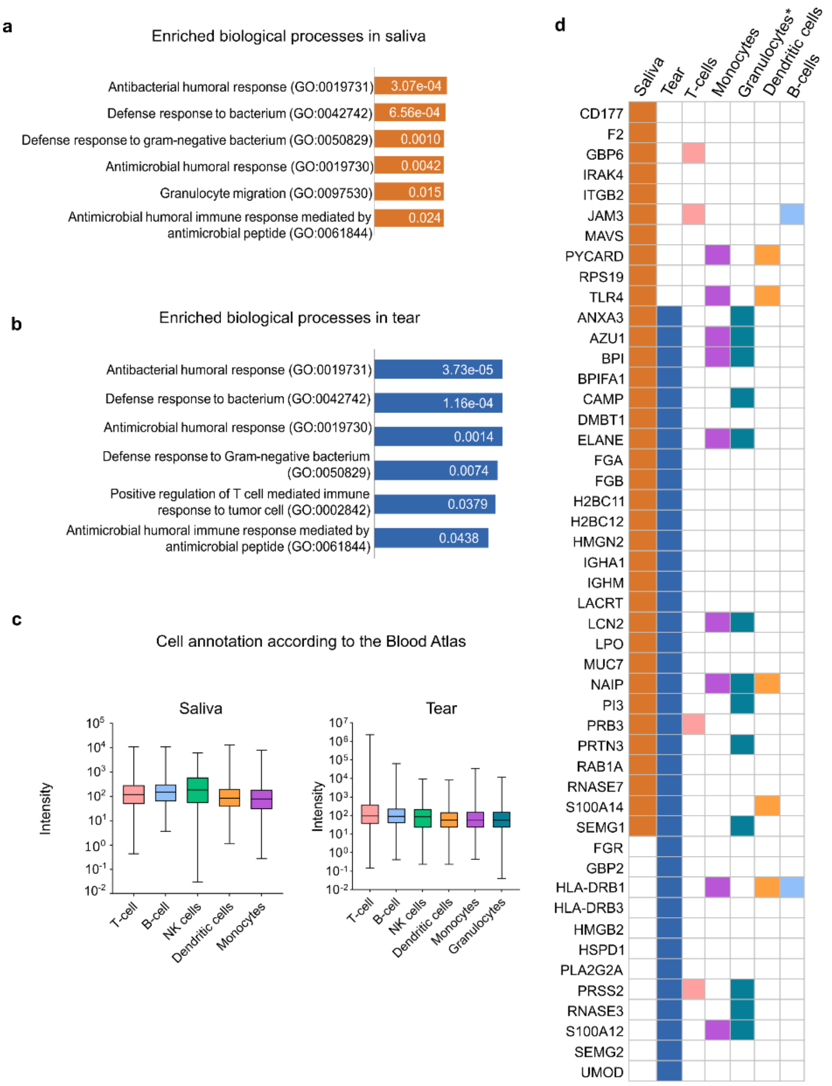


**Supplementary Figure 10. Enrichment analysis-driven cell bulk annotation.** Enriched biological processes in the A) saliva and B) tear fluid proteome indicates events of the immune response. C) Proteins involved in those biological processes were annotated using the single-cell data from The Blood Atlas to identify potential cell types present in the saliva and tear fluid. D) Potential cell types associated with the identified proteins in saliva and tear fluid annotated according to The Blood Atlas. *The mononuclear cell enrichment procedure was performed on saliva, and thus, the proteins associated with granulocytes were annotated exclusively for tear fluid.
